# Pathogenic IgG from long COVID patients with neurological sequelae triggers sensitive but not cognitive impairments upon transfer into mice

**DOI:** 10.1101/2025.11.20.689423

**Authors:** Margaux Mignolet, Catherine Deroux, Thomas Florkin, Valéry Bielarz, Kathleen De Swert, Nicolas Halloin, Lindsay Sprimont, Aurélie Ladang, Fabienne George, Jacques Gilloteaux, Laurence Abeloos, Johan Van Weyenbergh, Marc Jamoulle, Claire Diederich, Nicolas Albert Gillet, Pierre Bulpa, Charles Nicaise

## Abstract

Approximately 30% of long COVID patients still experience neurological symptoms (brain fog, pain, chronic fatigue) more than 4 months after the onset of COVID-19. This condition, known as ‘neurological long COVID’, remains poorly understood and might be explained by a persisting autoimmune response against nervous-derived self-antigens. The aim of this study is to determine whether IgG autoantibodies from long COVID patients with neurological sequelae can bind to central or peripheral nervous system epitopes and triggers neuropsychiatric symptoms upon passive transfer into mice, thereby mirroring patient-reported manifestations. Long COVID patients meeting the 2021 consensus WHO definition were included following a standardized neuropsychological assessment, while excluding patients with a medical history of autoimmune and neurological disorders. Age- and sex-matched asymptomatic individuals were used as healthy controls. Total IgGs were isolated using protein G purification and injected intraperitoneally into C57Bl6/J mice for four consecutive days. During the two weeks post-injections, behavioral tests assessed mechanical allodynia, thermal hyperalgesia, spatial working memory, depression, and anxiety. Mice injected with IgG from long COVID patients showed no difference with the control group in terms of anxiety or depression behaviors, as well as no impairment of short- or long-term spatial memories and thermal hyperalgesia. However, they displayed a transient decrease of paw withdrawal threshold during the first week. This effect was abolished when IgG-depleted serum or papain-digested IgGs were transferred. IgG from long COVID patients accumulated in the lumbar dorsal root ganglia of injected mice and colocalized with proprioceptive and nociceptive sensory neurons, without inducing local neuroinflammation or astrogliosis. These data demonstrate that IgGs from long COVID patients bind to peripheral sensory neurons and induce pain-related symptoms in mice. Our findings also support the hypothesis that autoantibodies mediate pain-related pathophysiology in the spectrum of long COVID symptoms.

## Introduction

Although patients with COVID-19 can recover completely, approximately 10% of SARS-CoV-2 infected people develop long-lasting symptoms after the acute phase of the disease [1]. This condition is termed as long COVID (LC) or post-acute sequelae of COVID (PASC). The U.S. National Academies of Sciences, Engineering and Medicine defines it as “*an infection-associated chronic condition that occurs after SARS-CoV-2 infection and is present for at least 3 months as a continuous, relapsing, remitting, or progressive disease state that affects one or more organ systems*” [2]. More than two hundred symptoms have been associated with LC, such as chronic fatigue, cognitive impairments and breathing issues. It is estimated that approximately 30% of LC patients suffer from cognitive impairment (brain fog, memory loss, concentration and attention deficits, anxiety, depression) [3–5] and 30% suffer from pain-related symptoms (myalgia, arthralgia, burning or prick sensations) [6–8]. Among these symptoms, 4-8% are estimated to experience neuropathic pain, typically characterized by allodynia or hyperalgesia [7, 8]. Other studies based on self-reported assessments of neuropathic symptoms suggest a prevalence up to 25% [9, 10]. So far, the underlying pathophysiology of such LC neurologic sequelae remains elusive, and no effective treatment is available for this disabling condition of daily life.

Given the broad heterogeneity of symptoms, several mechanisms have been proposed: SARS-CoV-2 viral persistence, chronic tissue inflammation, immune dysregulation, thrombo-inflammation, or latent viruses’ reactivation [11]. Among the immune dysregulation mechanisms, autoimmunity has been put forward as it is well described as part of the post-acute infection syndrome [12]. According to large cohort retrospective studies, the risk of developing new-onset autoimmune diseases significantly increases following acute COVID-19 [13–16]. Hence, we hypothesized that autoimmunity could similarly be triggered by an overstimulation of the immune system during acute SARS-CoV-2 infection and autoantibodies could arise following molecular mimicry, bystander activation or epitope spreading [17–20].

COVID-19 patients with neurological manifestations carry autoantibodies targeting nervous system antigens in serum or cerebrospinal fluid (e.g. APP, BDNF, mGluR5, NMDAR, myelin proteins) [21–25]. Anti-G protein-coupled receptor antibodies are frequently detected in LC patients with neurological symptoms [26, 27]. Moreover, months after acute infection, autoantibodies directed against nervous epitopes (e.g. myelin, NMDAR, GAD65) can still be detected in the serum and cerebrospinal fluid of LC patients with neurological impairment [28–31]. Interestingly, applying therapeutic plasmapheresis to LC patients significantly improved their clinical features and is associated with a reduction of autoantibody levels [32].

The aim of this study was to determine whether IgG antibodies from LC patients with neurological symptoms can bind to epitopes within the central and peripheral nervous systems and produce neuropsychiatric symptoms after passive transfer to mice. In order to directly implicate IgGs as the main drivers of these symptoms, the effects of purified patient IgG were compared with those of IgG-depleted serum and enzymatically-digested IgGs. We also investigated the specific cellular targets within the nervous system, aiming to bridge the gap between autoantibody presence and functional consequences. Following enrollment of LC patients with neurological symptoms and age- and gender-matched healthy control (HC) subjects, purified human IgGs were passively transferred into C57Bl/6J female mice. Along two weeks post-injection, pain-, cognitive-, anxiety- and depressive-like behaviors were followed-up in those mice.

## Materials and methods

### Patient selection

Human ethic project was approved by the CHU-UCL Namur ethics Committee (157.2022). Written and informed consent were obtained for all study participants. LC patients’ selection relied on the 2021 WHO consensus definition. Inclusion criteria for long COVID patients (LC; n=13) were: (i) 18 years of age or over; (ii) previous SARS-CoV-2 infection confirmed by a PCR or antigen test; (iii) presence of persistent symptoms (cognitive impairments, pain, fatigue) for at least 2 months that cannot be explained by an alternative diagnosis. Age- and sex-matched individuals (n=10) without morbidities served as healthy control group (HC) (Table 1). Their inclusion criteria were the same as above, except absence of persistent symptoms for at least 2 months since SARS-CoV-2 infection. Exclusion criteria for all participants were: (i) medical history of chronic pain, depression, cognitive impairments before SARS-CoV-2 infection; (ii) diagnosis of stroke, epilepsy, or neurodegenerative diseases (e.g. Alzheimer’s disease, amyotrophic lateral sclerosis, Parkison’s disease); (iii) diagnosis of autoimmune disease (e.g. rheumatoid arthritis, Sjogren syndrome, systemic lupus erythematosus, multiple sclerosis, type I diabetes). For all the subjects, the clinical interview about the relevant medical SARS-CoV-2 history (disease severity, time of infection, vaccination status) and persistent symptoms since SARS-CoV-2 infection (Table 1) was performed by an ICU COVID expert (PB) and a MD general practitioner (MJ). The neuropsychological assessment was performed by a trained neuropsychologist (CD) and included: Montreal Cognitive Assessment (MoCA), Symbol Digit Modalities Test (SDMT), STROOP test, D2 test, TAP Go/No go, TAP divided attention task, Beck Depression Inventory, Hospital Anxiety and Depression Scale (HADS) as well as a Numeric Pain Rating Scale (NPRS) and DN4 questionnaire (Table 2). All the neuropsychological tests were performed on the day of blood sampling. Serum was isolated within 30 minutes after blood sampling and kept at 4°C until IgG purification or aliquoted at - 20°C for the other protein assays.

**Table 1.** Patient demographics. Clinical data related to age, sex, SARS-CoV-2 infection(s), vaccination status, blood sampling and patient-reported symptoms are presented. P = percentiles. Statistical comparisons between HC and LC groups were performed using the Mann-Whitney U test or Fisher’s exact test, when appropriate.

|  |  | HC (n=10) | LC (n=13) | *p-value* |
| --- | --- | --- | --- | --- |
| **Age (years)** | Mean [range] | 48 [26-61] | 47.5 [28-64] | 0.7435 |
| **Sex** | Female | 8 | 12 | 1.0000 |
|  | Male | 2 | 1 |  |
| **Educational level** | • 1 | 0 | 1 | 1.0000 |
|  | • 2 | 0 | 0 |  |
|  | • 3 | 10 | 12 |  |
| **Hospital admission during acute COVID-19** | YES | 0 | 1 | 1.0000 |
|  | NO | 10 | 12 |  |
| **Date of infection with SARS-CoV-2** | • March 2020 to October 2020 | 0 | 6 | n.a. |
|  | • November 2020 to December 2021 | 2 | 6 |  |
|  | • January 2022 to December 2023 | 3 | 1 |  |
|  | • Unknown | 5 | 0 |  |
| **Number of infections with SARS-CoV-2** | Mean | 0.89 [0-2] | 1.9 [1-6] | 0.0617 |
| **COVID-19 vaccination status** | • Number of vaccination (mean) | 2.44 [0-4] | 3.4 [0-5] | 0.0438 |
|  | • Type of vaccine : |  |  |  |
|  | - AstraZeneca | 2 | 2 |  |
|  | - Moderna | 3 | 2 |  |
|  | - Pfizer | 6 | 10 |  |
|  | • Before sampling | 9 | 12 |  |
| **Time from first infection to sampling (months)** | Mean [range] | 25.25 [11-34] | 36.2 [22-44] | 0.0452 |
| **Patient-reported complaints at the time of sampling** | • Cognitive | 0 | 13 | <0.0001 |
|  | • Depression | 0 | 5 |  |
|  | • Pain | 0 | 13 |  |
|  | • Fatigue | 0 | 13 |  |

**Table 2.**
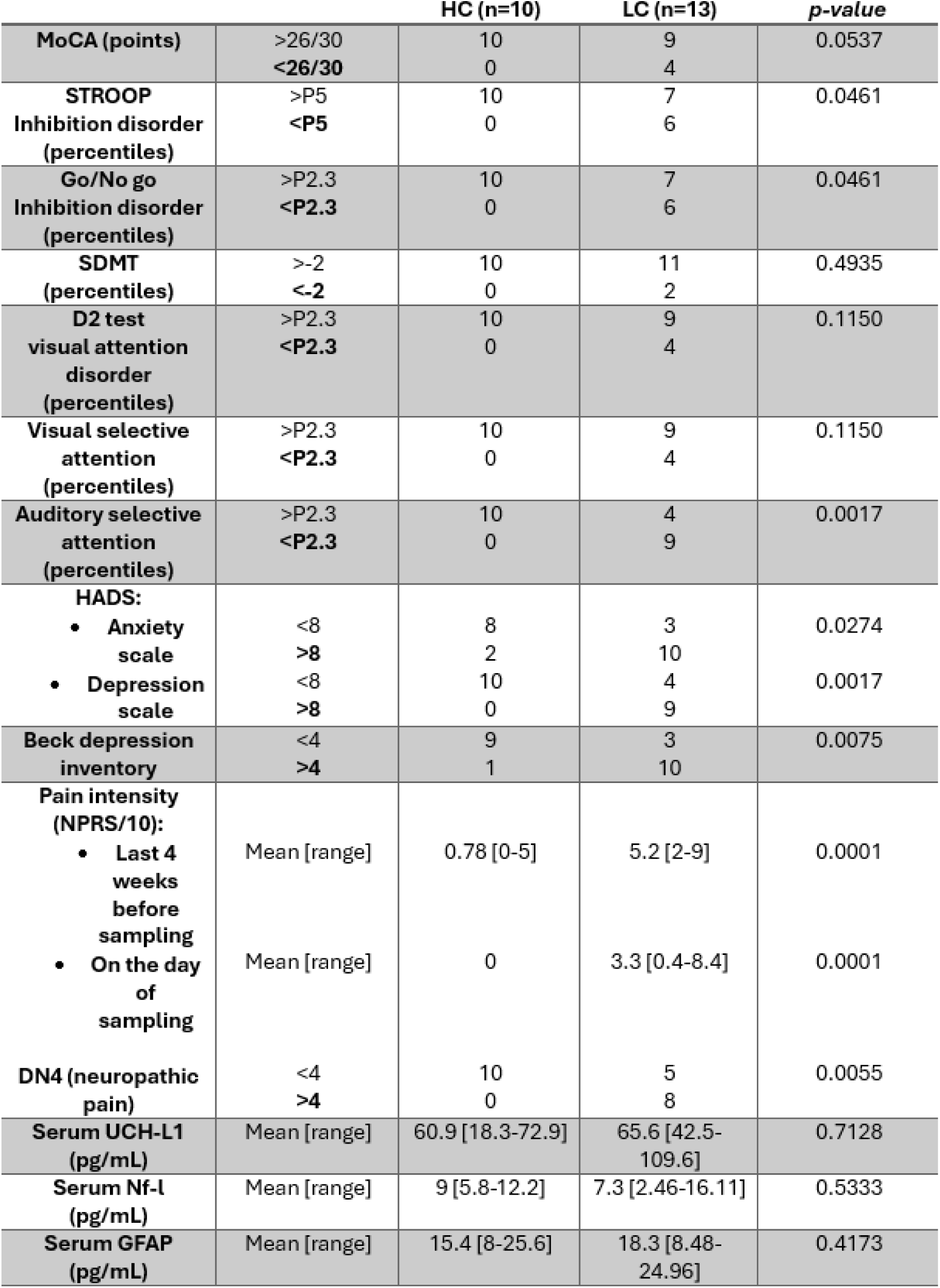
Neuropsychological assessment and biomarkers of neural damages. Expert-based cognitive testing included measures of memory (MoCA), executive functions (STROOP, Go/No Go task), attention (SDMT, D2 test, visual and auditory attention). Anxiety (HADS), depression (HADS, Beck) and pain intensity (NPRS, DN4) were assessed based on patient-reported outcomes using standardized questionnaires. Mean values for UCH-L1, Nf-L, GFAP serum levels are expressed in pg/mL. P = percentiles. Statistical comparisons between HC and LC groups were performed using the Mann-Whitney U test or Fisher’s exact test, when appropriate. Abbreviations: GFAP, glial fibrillary acidic protein; HADS, hospital anxiety and depression scale; MoCA, Montreal Cognitive Assessment; Nf-L, neurofilament-light; NPRS, numeric pain rating scale; SDMT, Symbol Digit Modalities Test; UCH-L1, ubiquitin carboxy-terminal hydrolase L1.

### IgG purification

For IgG purification from LC patients (n=13) or HC (n=10), serum was diluted 1:1 with glycine 0.1M NaCl 3M pH 8.9 and passed through a protein G column (Cytiva, Uppsala, Sweden) previously equilibrated with glycine 0.1M NaCl 3M pH 8.9. The column was rinsed with glycine 0.1M NaCl 3M pH 8.9, and the IgG-depleted sera was separately collected. The bound IgGs were eluted using glycine 0.1M pH 2.3 and immediately neutralized with Tris 1M pH 9.8. The column was then re-equilibrated with glycine 0.1M NaCl 3M pH 8.9. The eluate was dialyzed overnight at 4°C in PBS using a 12-14 kDa dialysis membrane (Spectrum Laboratories, Rancho Dominguez, CA, USA). The concentration of IgGs was determined using a Nanodrop 1000 (Thermo Scientific, Bleiswijk, The Netherlands). Finally, the IgG solution was stored at −20°C.

### Determination of Ig, UCH-L-1, Nf-l, and GFAP levels

Quantification of immunoglobulin isotypes (IgG, IgA, IgM) in sera and in purified IgG fractions was performed using Alinity 3 point-of-care diagnostic systems (AC01873, Abbott, Chicago, IL, USA). Quantification of IgE concentration was performed using human IgE ELISA kit (BMS2097, ThermoFisher Scientific, Waltham, MA, USA) according to manufacturer’s instructions. Neuronal and glial damage markers (Nf-l, UCH-L1, GFAP) were quantified in sera respectively using Quanterix SIMOA assay (Simoa® NF-light™ Advantage Kit (SR-X), Quanterix, Billerica, MA, USA) and Alinity 5 point-of-care diagnostic systems (AI01128, Abbott, Chicago, IL, USA).

### Papain enzymatic cleavage

Digestion buffer (PBS, 5 mM EDTA, 20 mM cysteine-HCl, pH 7.0) was prepared ex temporaneously. 2mL of immobilized papain (ThermoFisher Scientific, Waltham, MA, USA) were equilibrated by adding the digestion buffer and centrifuged at 1026 x g for 5 minutes to pellet the resin. This step was done twice. The IgG sample was then diluted 1:1 with the digestion buffer. Approximately 12mL of the solution were added to the immobilized papain so that the IgG solution was adjusted to 20mg/mL. Samples were incubated overnight at 37°C on a rotating platform. After the incubation period, the digested IgGs were separated from the immobilized papain by centrifugation (3912 x g for 5 minutes) to pellet the resin and collect the supernatant. The digested IgG solution was then stored at −20°C.

### Antigen microarray

Serum samples were screened for autoantibodies profiling using a commercial antigen array platform (Genecopoeia, Rockville, MD, USA). Briefly, sera were hybridized onto nitrocellulose filters adherent to glass slides to microarray slide spotted with 120 known autoantigens (PA002 OmicsArray^TM^ Brain and Central Nervous System Disorders Antigen Microarray). Slides were incubated with fluorescently coupled anti-IgG secondary antibodies and were imaged using a GenePix 4000B scanner. The Mapix software was used to analyze raw data (Innopsys France, Carbonne, France). Raw fluorescence data was normalized to PBS controls to get the Net Signal Intensity (NSI) of each antigen in each sample. The data presented in the heatmap are NSI normalized to internal Ig control batches according to manufacturer’s instruction.

### Western blotting

Serum and purified IgG samples were mixed with 5% β-mercaptoethanol and denatured via boiling for 5 minutes prior to gel loading. Serum or IgG (1 µg/well) were loaded on 10% polyacrylamide gel, separated via SDS-PAGE and transferred to a nitrocellulose blotting membrane (Protran, Amersham, Buckinghamshire, England). Membrane blocking was made using a bovine serum albumin 5% + TBS-Tween 0,1% solution for 1 hour at room temperature. Membranes were incubated with a solution of 5% TBST-BSA and anti-human IgG light and heavy chains antibody (1:50 000, 109-035-044, Jackson Laboratory, Bar Harbor, ME, USA) at room temperature for 1 hour. Prior to revelation, the membrane was rinsed 3 times with TBS-0.1% Tween and then incubated for 1 min in a chemiluminescent revelation solution (BM Chemiluminescence Blotting Substrate, Roche Diagnostics, Mannheim, Germany). The image acquisition was performed using an ImageQuant LAS 4000mini system (GE Healthcare, Little Chalfont, UK).

### Mice handling

The experimental procedure was approved by the Animal Ethics Committee of the University of Namur (ethic project UN 23-392 and UN 24-438). Mice were housed in a temperature-controlled environment with a 12-hour light/12-hour dark cycle. They had access to food and water *ad libitum*. Behavioral experiments were performed on female C57BL/6J mice (8-10 weeks old) purchased from Charles River Laboratories (Beerse, Belgium). Mice received intraperitoneally the human IgG solution (8mg/day) for four consecutive days. Each patient IgG batch was injected into one cohort of mice (n=10). Along two weeks post-injections, mice were submitted to behavioral tests (Fig.1A).

**Figure 1.**
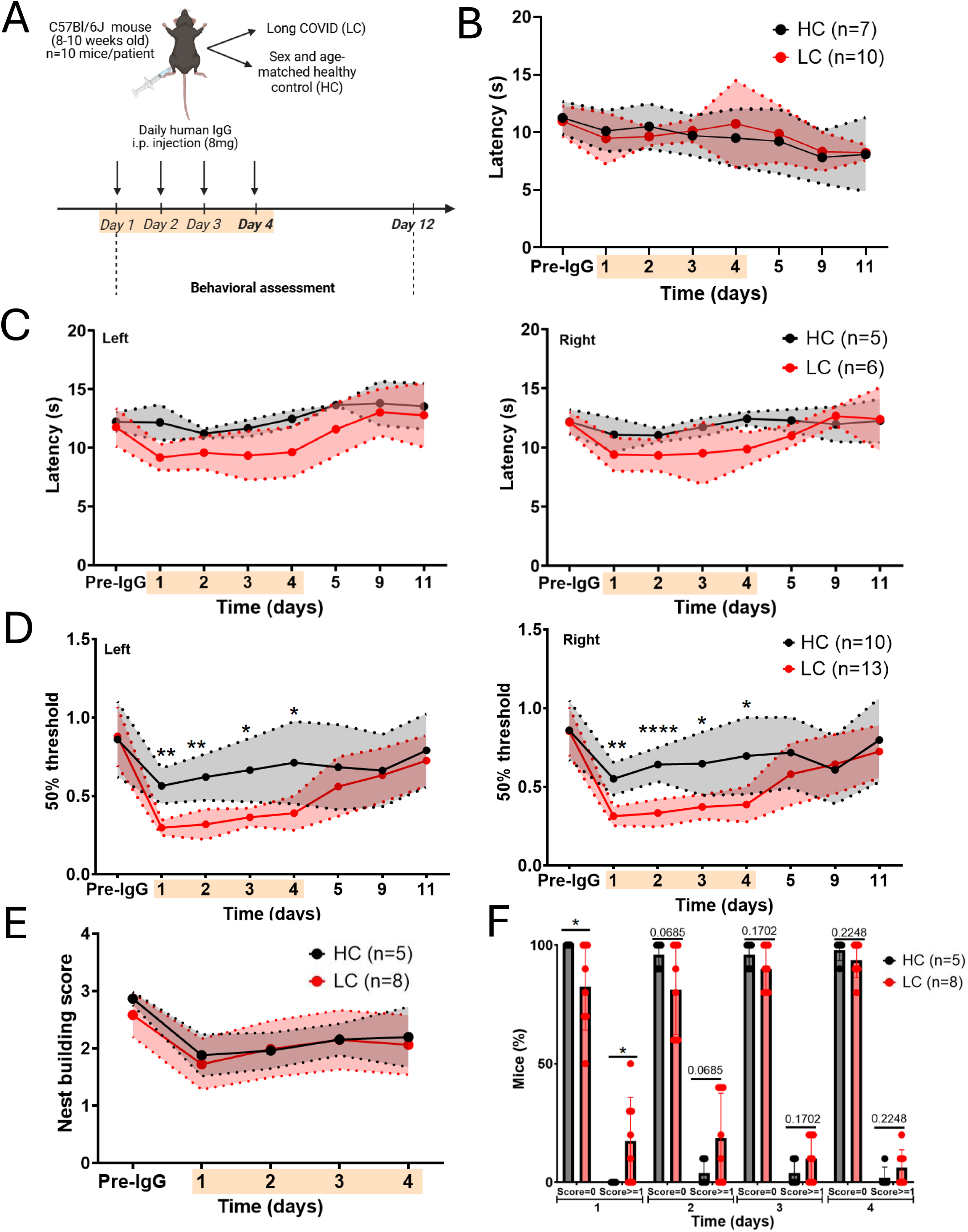
Pain-related behavioral tests in mice transferred with IgG from long COVID patients (LC) or healthy controls (HC). (A) Experimental timeline. 10 mice were used per patient IgG batch. n=5-10 HC and n=6-13 LC. The orange overlay on each timeline corresponds to the injection period. (B) Paw withdrawal latency at the hot plate test was unchanged between HC and LC groups. (C) Hind paw (left and right) withdrawal latency at the Hargreaves test did not differ significantly between HC and LC groups. (D) Hind paw (left and right) withdrawal threshold at the Von Frey filaments was significantly decreased during the first four days post-injection in LC condition. (E) The general well-being and motivation behavior were similar between groups of mice at the nest building score. (F) The percentage of mice with an abnormal Facial Grimace Scale (score ≥1) was significantly increased the first day post-injection in LC group. Mean ± SD. Mixed-effects model followed by a Holm-Sidak multiple comparison test between HC and LC *(*p<0.05, **p<0.01, ****p<0.0001)*.

A second experimental paradigm was conducted using a modified protocol. Mice received intraperitoneally either IgG-depleted serum, papain-digested IgG, or native IgG (8mg/injection) from LC patients. Pain-related behavioral tests were performed as previously but over a period of two days (Fig.2A). Mice were randomized between cages, and the experimenter was blinded to their treatment.

**Figure 2.**
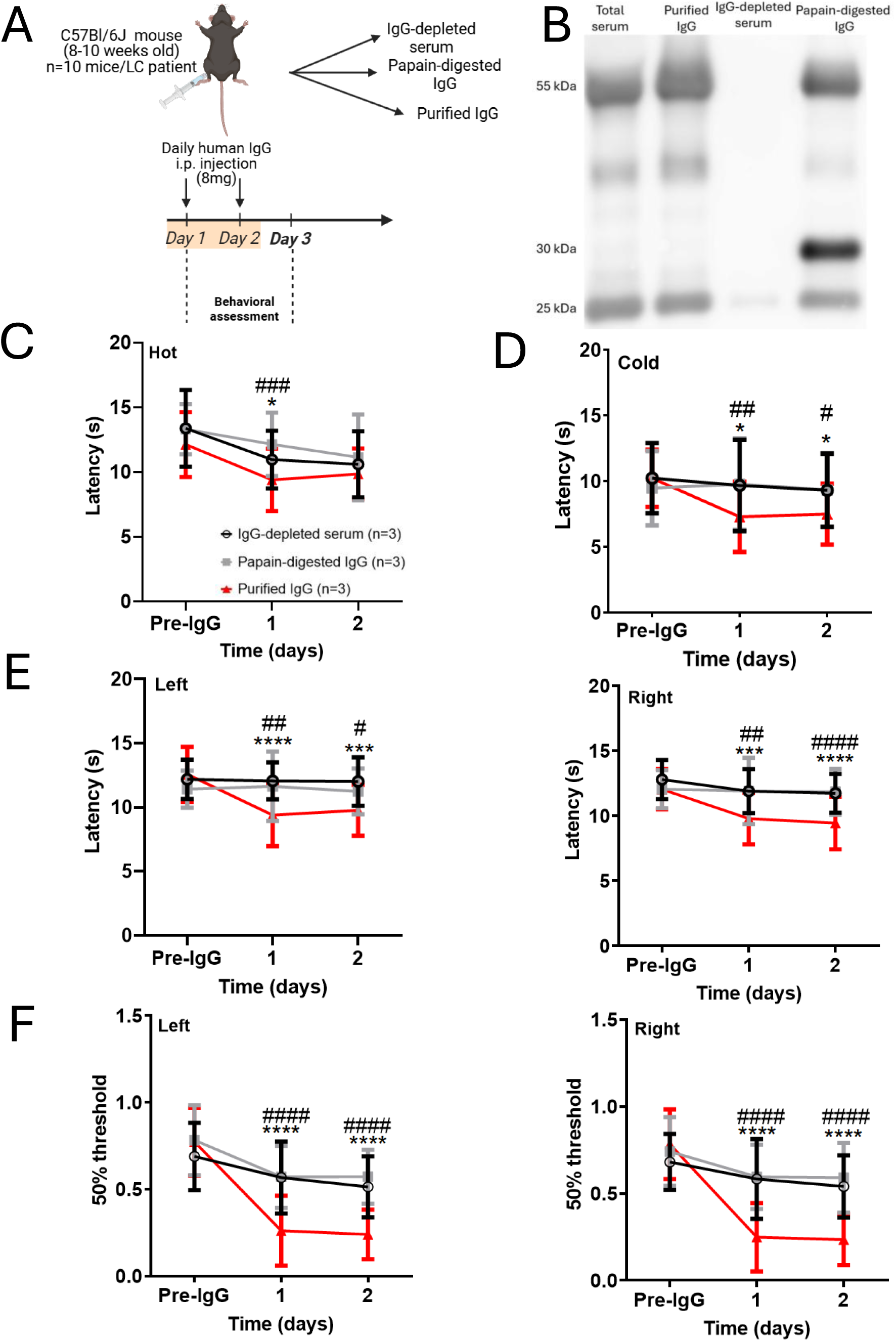
Pain-related behavioral tests in mice transferred with purified IgG, papain-digested IgG or IgG-depleted serum, from long COVID patients. (A) Experimental timeline. 10 mice were used per patient IgG batch, n=3 LC IgG batch. (B) The abundances of heavy (55 kDa), light chains (25 kDa), Fc (30 kDa) and Fab fragments (25 kDa) were assessed in total human serum, purified IgG fraction, IgG-depleted serum and papain-digested IgG fraction by immunoblotting. The presence of an extra band at 30 kDa confirmed the partial digestion of IgG heavy chains after papain incubation. (C) Paw withdrawal latency at the hot plate test was rescued the first day post-injection in mice receiving depleted serum and papain-digested IgG. (D) Paw withdrawal latency at the cold plate test was fully rescued in mice receiving depleted serum and papain-digested IgG compared to native purified IgG. (E) Hind paw (left and right) withdrawal latency at the Hargreaves test were significantly different in mice receiving depleted serum and papain-digested IgG compared to native purified IgG. (F) Hind paw (left and right) withdrawal threshold at the Von Frey filaments was fully rescued in mice receiving depleted serum and papain-digested IgG. Mean ± SD. Mixed effects model followed by a Holm-Sidak multiple comparison test between purified IgG and the IgG-depleted serum group *(*p<0.05, ***p<0.001, ****p<0.0001)* and between purified IgG and the papain-digested IgG group *(^#^p<0.05,^##^p<0.01, ^###^p<0.001, ^####^p<0.0001)*.

### Behavioral testing

Before any tests, mice were kept in their cages to acclimatize (10-15 minutes) to the experimental room. Baseline measurements were performed before any IgG injection.

#### Hot and cold plates

The hot plate analgesia meter (Columbus Instruments, Columbus, OH, USA) was set to 52°C while the cold plate system (ElectraCOOL™ TCP50™, Advanced thermoelectric, Melbourne, FL, USA) was set to 4°C. Mice were placed onto each platform surrounded by transparent Plexiglas walls. The time the mice took to show signs of thermal distress (paw licking, jumping, hind paws stomping) was measured. If mice did not show any of these signs, they were removed from the plate after 30 seconds to prevent tissue damage.

#### Hargreaves test

Mice were placed onto a glass surface, separated by a Plexiglass wall, and allowed to acclimate for 10 minutes. To observe a variation in thermal sensitivity among mice, a radiant thermal beam was placed under their left and right hind paws to provoke a withdrawal response using a Plantar Test Analgesia Meter (Model 390, IITC Life Science, Los Angeles, CA, USA). The time separating the stimulus onset and the paw withdrawal, called latency (in seconds), was recorded. If mice did not show signs of thermal distress (licking of the paw, paw withdrawal), the beam was removed after 20 seconds to prevent tissue damage. Each hind paw was stimulated three times in total with five minutes between each stimulation. The latencies of each individual were averaged.

#### Von Frey filaments

Von Frey filaments were calibrated monofilaments (0.008-2g) used to apply a specific pressure. Mice were tested in individual cages equipped with a stainless-steel wire meal that allowed full access to the paws (Bio-VF-M, Bioseb, Vitrolles, France). Stimulation with the filaments was limited to the medio-plantar area of the paw. One of a series of 8 Von Frey filaments, with logarithmically increasing stiffness, were applied to the plantar area for 2 to 3 seconds. The force applied should be sufficient to induce a slight buckling against the paw. Immediate licking of the filament stimulation area was considered a positive response to filament stimulation. The Chaplan’s Up-and-Down method was used [33], meaning that the test was initiated with the 0.4g hair and then the stimuli was presented consecutively, up or down. If there was no paw withdrawal response to the initial stimulus, a stronger stimulus was presented. If a paw withdrawal response was observed with the initial stimulus, a weaker stimulus was presented. This was done until the absence of paw withdrawal (if descending) or the presence of paw withdrawal (if ascending).

#### Nest Building Score

Mice were isolated in a new cage containing new nesting material. 7 hours post-IgG injection, nest construction was blindly evaluated by the experimenter using a rating scale from 1 to 3 (1 = no nest construction; 2 = partial nest with scattered material, 3 = perfectly constructed nest). This evaluation took place every day during the 4 days of IgG injection, and each day the mice were isolated in a cage with new nesting material. At the end of the 4-day injection period, the mice that were initially together in the same cage were again put back together in a new cage containing a mixture of the new nest and each other’s nests.

#### Facial Grimace Scale

Videos of mice were recorded every day during the first week of IgG injection. They were then blindly evaluated by the experimenter and scored (0 = not present; 1 = moderately present; 2 = obviously present). Five facial expressions were analyzed: orbital tightening, nose bulge, cheek bulge, ear position, and whisker change.

#### Barnes maze

Briefly, the mouse was dropped in the center of an elevated circular platform (122 cm diameter) with 40 evenly spaced holes (5 cm diameter). An escape box (22,5c m length x 8,5 cm wide × 10,5 cm high) was located underneath one hole. This box was maintained at a fixed location for the whole duration of the test. Four visual cues were located at regular intervals around the table so that mice can use them to orient themselves. This test was done in three steps: *Habituation phase* (day 1) - mice were placed in the box for 60 seconds and then in the center of the table. They were allowed to explore it until they entered the escape box, or 300 seconds had elapsed. *Acquisition training* (day 1-5) – it started at least 1-hour after the habituation phase. Three trainings each day for five days were performed during which the animal was placed in the center of the maze in a covert start box for 8 seconds and was then allowed to explore the maze for 180 seconds. If by the end the mouse had not entered the escape box, the mouse was gently guided towards the corresponding hole and allowed to remain there for 60 seconds. *Post-injection measures* - during the two weeks post-injection, mice only performed one trial at specific time points. Between each trial, the maze was thoroughly cleaned with ethanol to remove olfactory cues. The primary latency (time to locate the target hole) and total latency (time to enter the escape box) were recorded.

#### Y-maze

The Y-maze (arm’s length 40 cm x 8 cm wide, center zone 8 cm diameter) was made of three symmetrical arms (A, B, C). The mouse was placed at the same end of one arm of the Y-shaped maze and allowed to explore it for 300 seconds. The total distance traveled was determined by a tracking software (EthoVision XT 18, Noldus System, Netherlands) and served as a measure of fatigue. The number of alternations between each arm (arm A, arm B, arm C) and the number of entries in each arm were monitored by a video tracking system (EthoVision XT 18, Noldus System, The Netherlands). The percentage of alternations was then calculated as follows: (total number of alternations/number of arms entered) × 100.

#### Elevated-plus maze

Elevated-plus maze consisted of a plus-shaped maze elevated above the ground (51 cm) with two opposite closed arms (30 cm length x 5 cm wide), two opposite open arms (30 cm length x 5 cm wide) and a central square (5cm diameter). Mice were placed at the center of the maze, head facing an open arm, and explored the maze for 300 seconds. The time spent in open and closed arms was measured by a video tracking system (EthoVision XT 18, Noldus System, The Netherlands).

#### Light/Dark box

The apparatus consisted of a box (50 cm length x 24 cm wide x 24cm high) equally divided into a bright and a dark compartment. An opening of 7 cm high and 7 cm wide connects the two parts. Mice were placed in the bright chamber and explored the box for 300 seconds. The time spent in dark and bright chambers was measured by a video tracking system (EthoVision XT 18, Noldus System, The Netherlands).

#### Tail suspension test

The test consisted of a tail suspension apparatus (42 cm length x 14 cm wide x 24,5cm high) with three separate compartments so that mice cannot observe and interact with each other. A piece of tape was placed on the tail of the mouse, precisely 2 cm from its tip. The tail itself was attached to a hook in the middle of the suspension cage. For 360 seconds, the duration of the movements of the mouse suspended by the tail (periods of agitation and periods of immobility) were recorded.

### Euthanasia and histology

For euthanasia, mice were anesthetized using Ketamine/Xylazine cocktail (120 mg/kg ketamine (Nimatek); 8 mg/kg xylazine (Sedaxylan)), perfused intra-cardially with cold 0,9% NaCl. The brain, spinal cord, and lumbar dorsal root ganglia (DRG) were freshly harvested, snap-frozen or fixed with 4% paraformaldehyde for histology analysis. Brain and spinal cord samples were dehydrated with a Leica HistoCore (Leica Biosystems, Nanterre, France) and embedded in paraffin. Ten µm-thick sections were obtained using a microtome (RM2145, Leica Biosystems, Nanterre, France). DRGs were washed with PBS and transferred in 30% sucrose solution at 4°C for 48-72 hours. Then, they were embedded in OCT compound, frozen in cold isopentane and stored at −80°C. Cryosections, 10µm in thickness, were obtained using a cryostat (CM1950, Leica Biosystems, Wetzlar, Germany).

#### Immunohistochemistry

Paraffin sections were dewaxed and sequentially rehydrated. Heat-induced epitope retrieval was performed using a 0.01M citrate buffer pH 6 in a 96°C water bath for 10 minutes. Following endogenous peroxidase blocking with 3% hydrogen peroxide for 10 minutes, sections were incubated with 5% goat serum-TBS for 15 minutes at room temperature, and were incubated overnight at 4°C with primary antibodies (listed in Table 3) or 4 hours at room temperature with anti-human IgG antibody diluted in 1% goat serum-TBS. Sections were then incubated with secondary antibodies (Vectastain ABC-HRP kit, Peroxidase [Mouse IgG] PK-4002 or [Rabbit IgG] PK-4001 or [Goat IgG] PK-4005, Vector laboratories, Newark, CA, USA) for 30 minutes at RT. Immunolabeling was revealed using 3,3’-diaminobenzidine (DAB, K3468, Dako, Santa Clara, CA, USA). Finally, the sections were counterstained with hematoxylin and mounted with DPX medium. Observations were obtained following slide scanning using Pannoramic Flash Desk DX digital scanner (3DHistech, Budapest, Hungary).

**Table 3.**
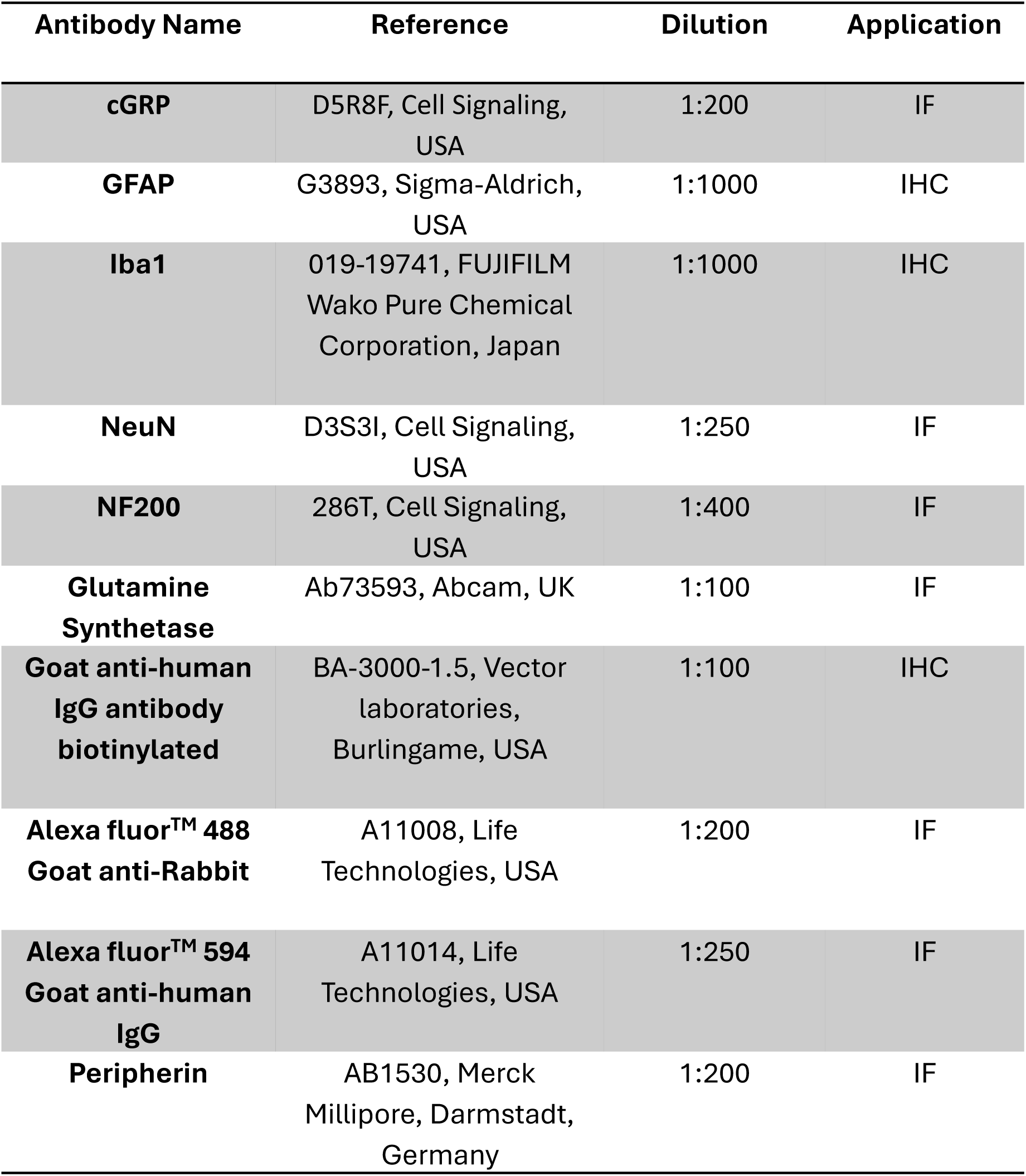
Primary and secondary antibodies for immunohistochemistry (IHC) or immunofluorescence (IF).

#### Immunofluorescence

Cryosections were washed with TBS and saturated with TBS-BSA 0.2% - Tween 0.02% for 1-hour at RT. Primary antibodies (Table 3) were diluted in the saturation solution and incubated on sections at 4°C O/N. After washing with TBS-Tween 0.02%, sections were incubated 1h at room temperature with secondary antibodies (Table 3). Nuclei were counterstained with Hoechst (1:200; 94403, Sigma, St Louis, MO, USA). Sections were mounted with Mowiol and kept at 4°C until imaging with LSM 900 confocal microscope (Zeiss, Germany). Images analyses were performed using the software ImageJ.

### RNA extraction and Quantitative PCR

Brain and DRG tissues were resuspended in 1 mL of TriZol reagent (Life Technologies, Bleiswijk, The Netherlands). Total RNAs were isolated according to manufacturer’s instructions (High Pure RNA Tissue kit 12033674001, Roche, Mannheim, Germany). RNA yield and purity were determined using a spectrophotometer NanoDrop 1000 (Thermo Scientific, Bleiswijk, The Netherlands). Total RNAs were reverse transcribed using the Super Script III Reverse Transcriptase kit according to manufacturer’s instructions (Invitrogen, Merelbeke, Belgium). cDNA samples were used to amplify genes of interest with Takyon SYBR Green kit (Eurogentec, Liège, Belgium) in a Light Cycler 96 device (Roche Diagnostics, Mannheim, Germany). Primer sequences (Eurogentec, Liège, Belgium) are as follow: *Gfap forward* 5’-GCCACCAGTAACATGCAAGA-3’; *Gfap reverse* 5’-CGGCGATAGTCGTTAGCTTC-3’; *Iba1 forward* 5’-CTTGAAGCGAATGCTGGAGAA-3’; *Iba1 reverse* 5’-GGCAGCTCGGAGATAGCTTT-3’; *Hprt forward* 5′-TGACACTGGCAAAACAATGCA-3′; *Hprt reverse* 5′-GGTCCTTTTCACCAGCAAGCT-3′. The relative gene expression was calculated using the ΔΔCq method with *hprt* as housekeeping gene.

### Statistical analysis

Data are presented as mean ± SD and the number of patients included are indicated by *n*. 10 mice were systematically used per patient IgG batch. Paired t-test (Suppl. Fig. 1), Mann-Whitney U test (Figs. 4, 6B, tables 1, 2), Welch’s t-test (Fig.8), Fischer’s exact test (Tables 1, 2), Kruskal-Wallis H test (Fig. 7B), and mixed model followed by Holm-Šídák multiple comparison test (Figs. 1B-F, 2C-F, 3A-F, Suppl. Figs. 2A-E, 3A-E.) were used to determine statistical significance (set at *p* < 0.05). Statistical analyses were performed on the software GraphPad Prism (v.10.2.3, La Jolla, USA) or in the R environment (v.2025.05.1+513).

**Figure 3.**
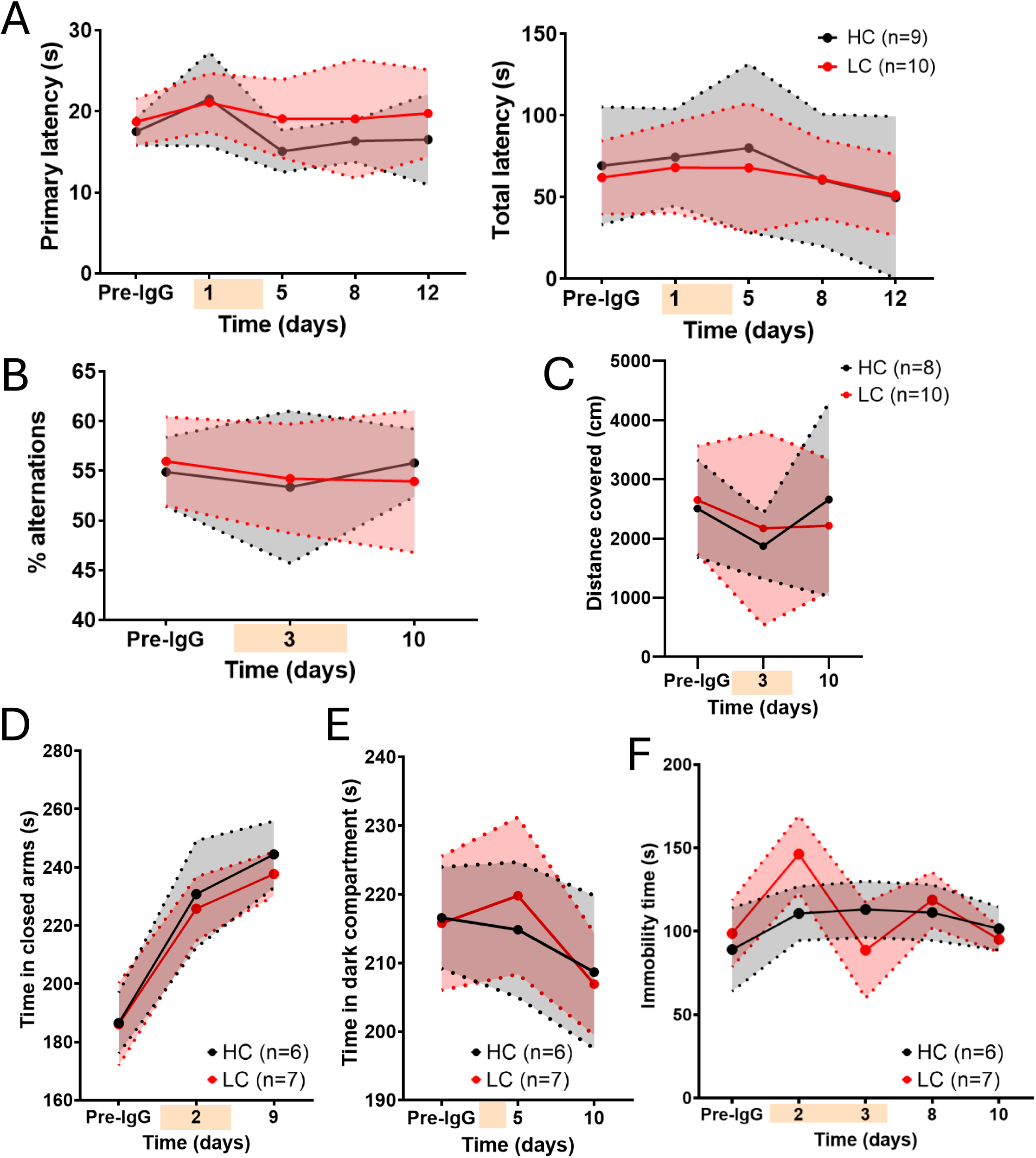
Spatial memory-, anxiety- and depression-related behavioral tests in mice transferred with IgG from long COVID patients (LC) or healthy controls (HC). 10 mice were used per patient IgG batch. n=6-9 HC and n=7-10 LC patients. The orange overlay corresponds to the injection period. (A) Primary and total latencies at the Barnes maze did not differ between HC and LC groups. (B) Alternation between the arms of the Y-maze did not differ between HC and LC groups. (C) Total distance traveled in the Y-maze reflecting the general locomotor activity was unchanged between experimental conditions. (D) Time spent in the closed arms of the elevated-plus maze, as a measure of anxiety, was similar between groups. (E) Time spent in the dark compartment of the light and dark box, as a measure of anxiety, was similar between groups. (F) Immobility time at the tail suspension test, as a proxy of depressive-like behavior in mice, did not differ between HC and LC groups. Mean ± SD. Mixed effects model followed by a Holm-Sidak multiple comparison test between HC and LC.

**Figure 4.**
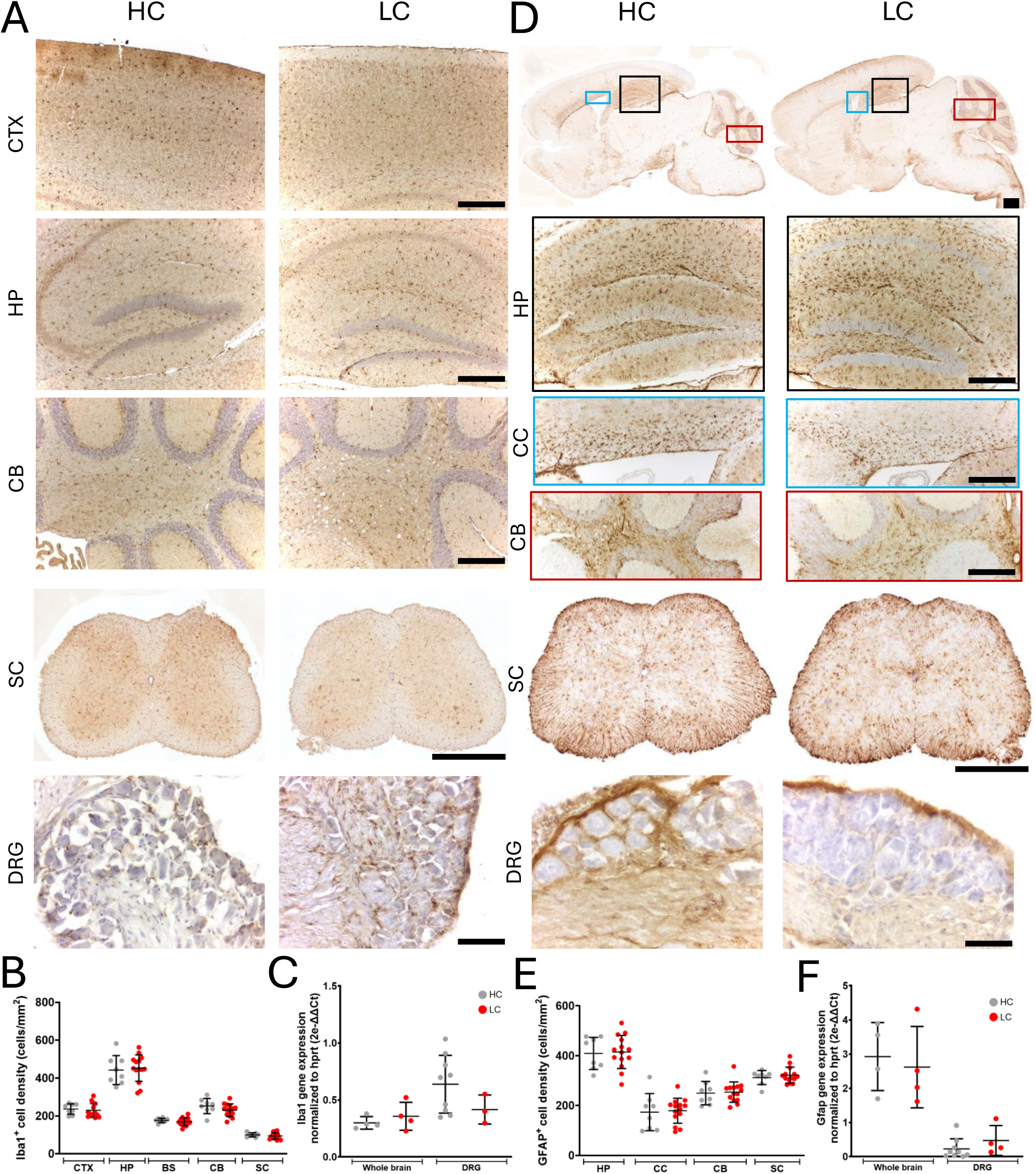
Assessment of glial activation in the central and peripheral nervous systems of mice transferred with human IgG. (A) Immunolabeling of Iba1 protein in the brain, spinal cord and DRG of mice. (B) The quantification of Iba1^+^ microglia density did not show any significant difference in the (sub)regions-of-interest between HC and LC groups. (C) The normalized expression of Iba1 mRNA did not show any significant difference in the whole-brain or DRGs between HC and LC groups. (D) GFAP protein was immunolabeled in the brain, spinal cord and DRG of mice. Scale bar represents 500 µm for brain and spinal cord and 100 µm for DRGs. (E) The quantification of GFAP^+^ astrocyte density did not show any significant difference in the (sub)regions-of-interest between HC and LC groups. (F) The normalized expression of GFAP mRNA did not show any significant difference in the whole-brain or DRGs between HC and LC groups. Mean ± SD. Statistical analyses were computed using a Mann-Whitney U test. *p-value=ns*. Abbreviations: CTX, cortex; HP, hippocampus; BS, brainstem; CC, corpus callosum; CB, cerebellum; SC, spinal cord; DRG, Dorsal Root Ganglia.

**Figure 5.**
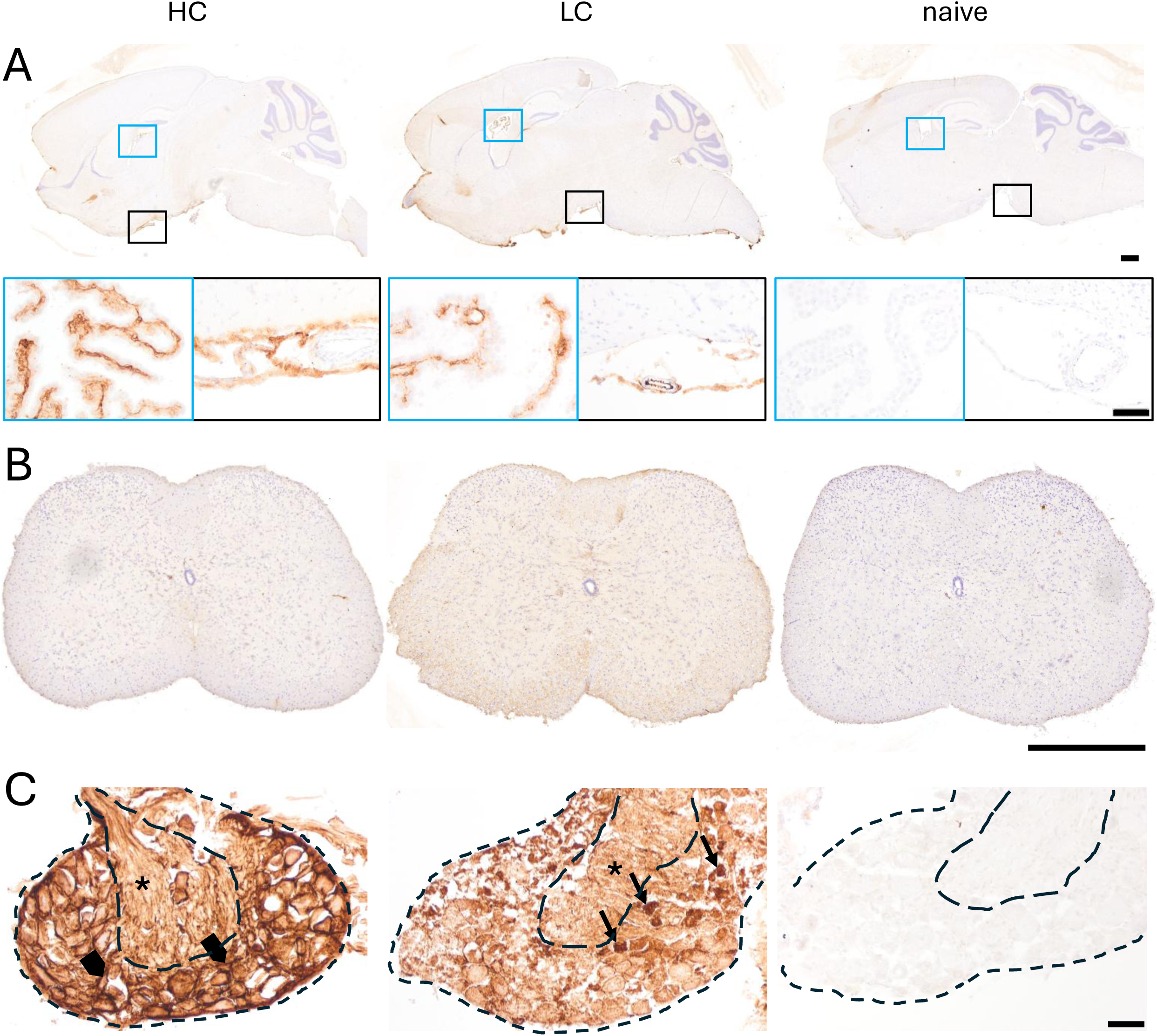
Distribution of transferred human IgG throughout the central and peripheral nervous systems of mice. (A) In brains, human IgG immunoreactivity was visualized in the connective tissue of choroid plexuses (blue square) and in the meningeal layer (black square), while not detected in the parenchyma, for both HC and LC conditions. No immunoreactivity was seen in the brains of naïve mice that were not transferred with human IgG. Scale bar represents 500 µm. Insets scale bar represents 50 µm. (B) In lumbar spinal cords, no immunoreactivity was observed for both HC and LC conditions, as well as naïve mice. Scale bar equals 500 µm. (C) In lumbar DRG, human IgG immunoreactivity was observed in the fiber-rich area (black star) and in the peri-neuronal spaces (arrowheads) in HC condition. Distribution of human IgG was strikingly different in LC condition, with immunoreactivity on sensory neuron cell bodies (arrows). No immunoreactivity was seen in the DRG of naïve mice. Scale bar represents 50 µm.

**Figure 6.**
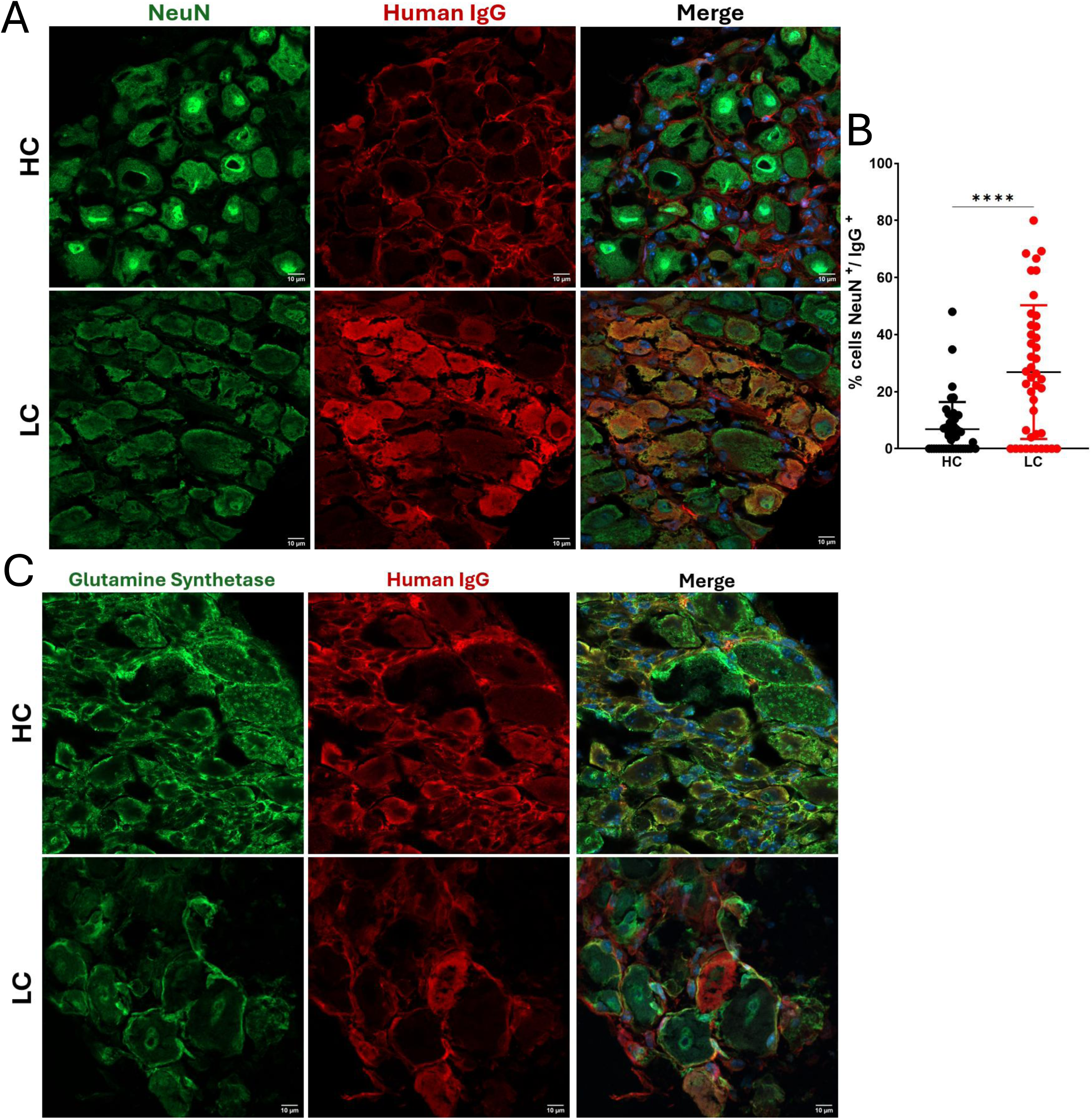
Tissue distribution of IgG from long COVID patients (LC) or healthy controls (HC) in the mouse DRGs. Human IgG were immunolabeled using fluorescent anti-human IgG antibody (red). Nuclei were counterstained with Hoechst (blue). (A) Neuronal nuclei and cell bodies in the murine DRG were co-labeled using fluorescent anti-NeuN antibody (green). (B) Quantification of NeuN^+^/IgG^+^ cells was performed using FIJI ImageJ and revealed a significant increase of double immunolabeled cells in LC condition compared to HC group. Each dot represents a colocalization. DRGs included in the analysis were representative from 4 HC condition (n=8 mice) and 3 LC condition (n=5 mice). Mean ± SD. Statistical analyses were performed using a Mann-Whitney U test *(****p<0.0001).* (C) Satellite glial cells were immunostained using fluorescent anti-glutamine synthetase antibody (green). No colocalization with human IgG was ever evidenced with glutamine synthetase. Scale bar represents 10 µm.

**Figure 7.**
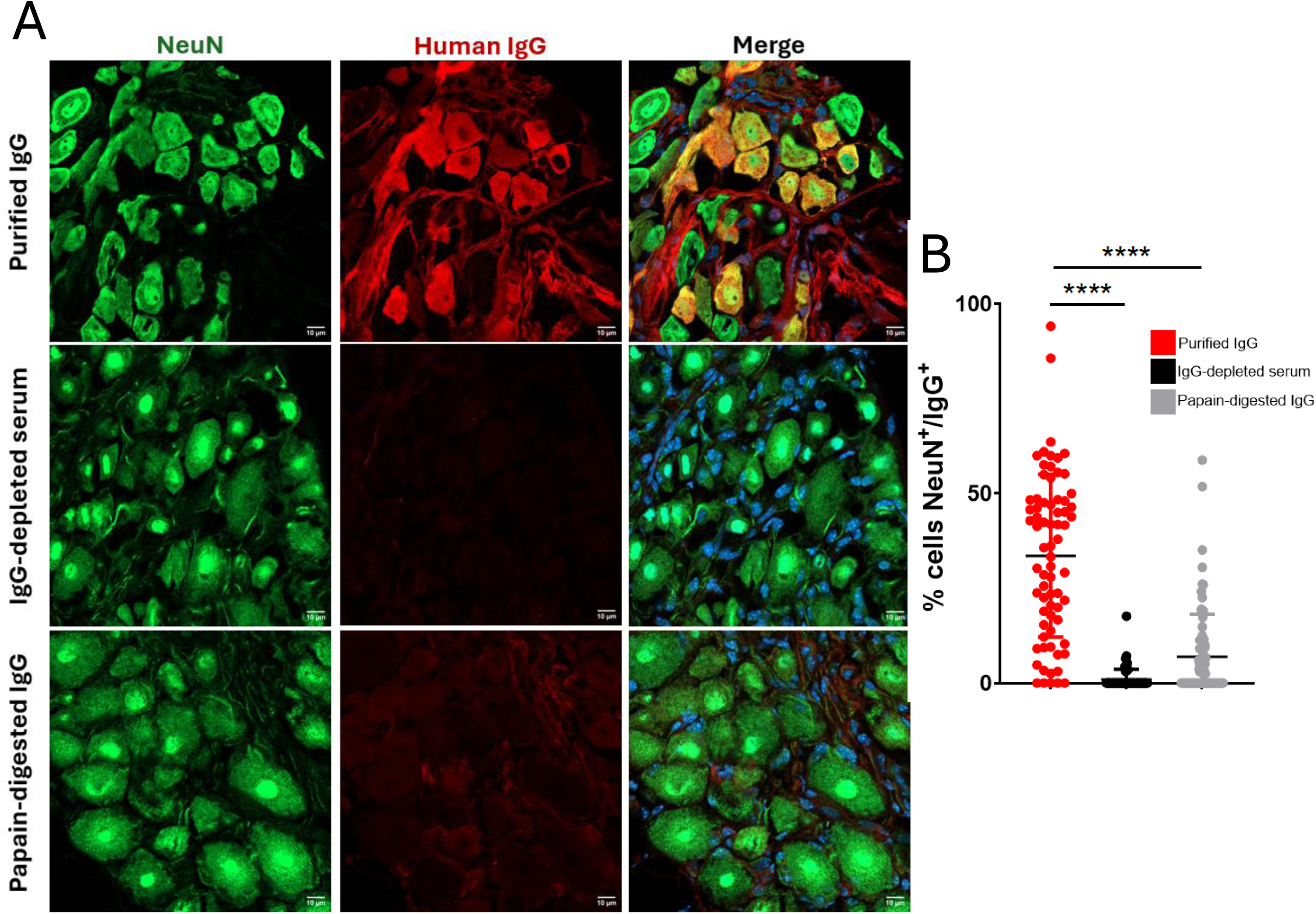
Detection of human IgG in the mouse DRG after transfer of native IgG, depleted serum, or papain-digested IgG from long COVID patients. Human IgG were immunolabeled using fluorescent anti-human IgG antibody (red). Nuclei were counterstained with Hoechst (blue). (A) Neuronal nuclei and cell bodies in the murine DRG were co-labeled using fluorescent anti-NeuN antibody (green). (B) Quantification of NeuN^+^/IgG^+^ cells was performed using FIJI ImageJ. The transfer of IgG-depleted serum or papain-digested IgG significantly abolishes their detection in the mouse DRG. Each dot represents a colocalization. DRG included in the quantitative analysis were from matched LC IgG-depleted serum (n=12 mice), papain-digested IgG (n=13 mice), and purified IgG (n=10 mice). Mean ± SD. Statistical analyses were performed using a Kruskal-Wallis comparison test *(****p<0.0001).* Scale bar represents 10 µm.

**Figure 8.**
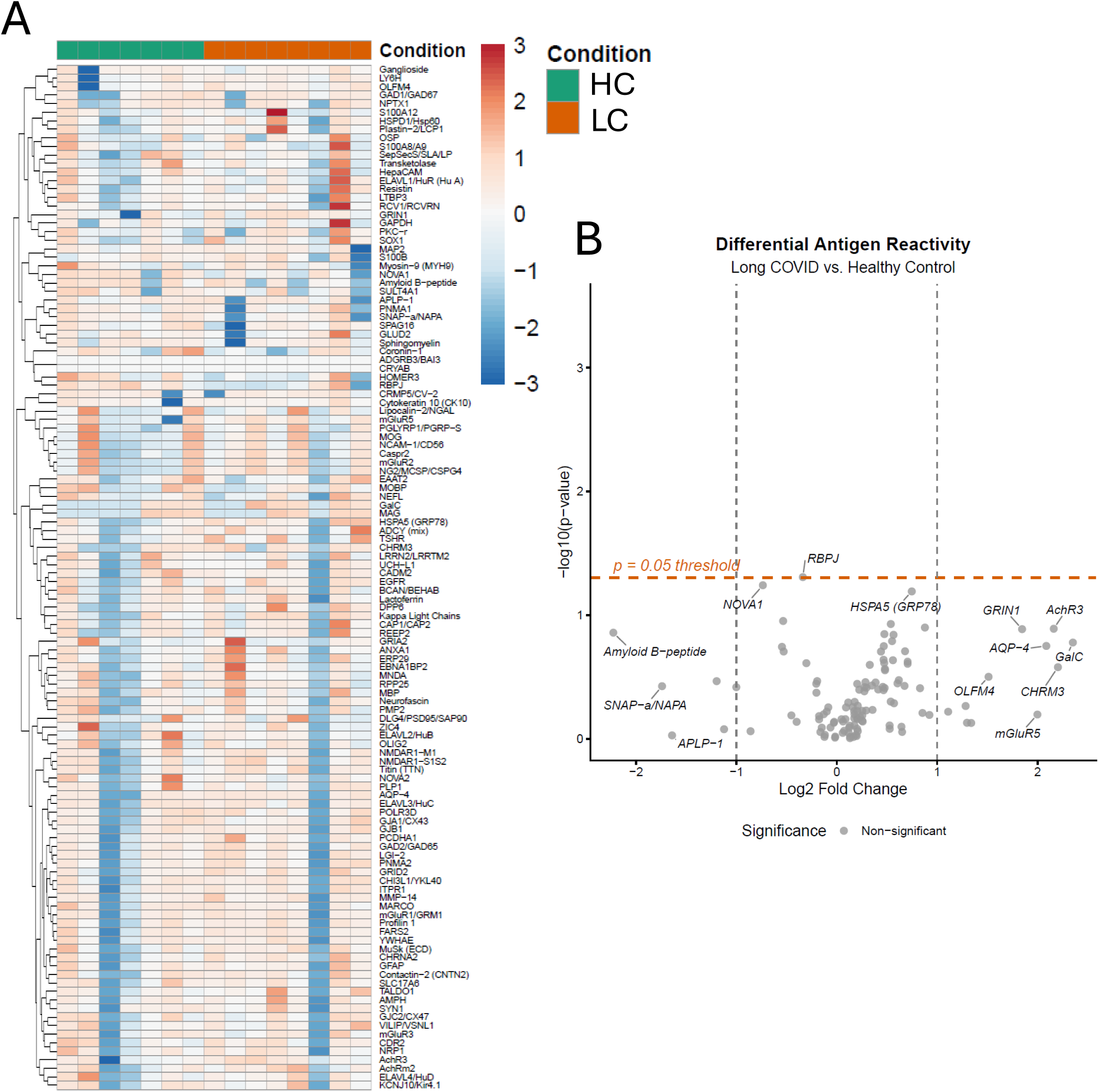
Levels of autoantibodies associated with known neurological disorders in the serum used for IgG isolation and transfer. (A) Heatmap displaying the antibody reactivity profile for 120 analyzed antigens in HC (n=7) and LC (n=8) patients. The color of each cell represents the row-wise Z-score, indicating the number of standard deviations a patient’s reactivity is from the mean of all patients for that specific antigen. Antigens are organized by hierarchical clustering to group those with similar profiles. (B) Volcano plot showing in x-axis the log Fold Change and in y-axis the log of p-value. Statistical comparisons were performed using a Welch’s t-test (p>0.05).

## Results

### Patients’ demographics and neuropsychological assessment

A total of 25 LC patients were screened and 13 were found eligible and enrolled in the study. The reasons for non-enrollment were lack of neurologic symptoms at the time of recruitment, or pre-diagnosed autoimmune or neurologic diseases. Clinical demographics and neuropsychological test performances for HC and LC patients are presented in Table 1 and Table 2. Our LC cohort included mostly women (M:F ratio 1:12), with a mean age of 47.5 years. Average time from the first SARS-CoV-2 infection to recruitment was 36.2 ± 6.9 months. All participants of the LC cohort (n=13) reported cognitive impairment, pain, and fatigue at the time of recruitment and blood sampling. Five LC patients also reported symptoms of depression, while none of the HC participants (n=10) reported such complaints (Table 1). Other clinical-relevant features included the patient-reported number of infections, the dates of infection, the vaccination status or type of vaccines (Table 1).

Expert-based neuropsychological assessment revealed that 6 LC patients exhibited inhibition disorders, 4 had visual attention disorders, and 9 showed signs of auditory attention disorders. Most of these LC patients presented symptoms of depression (n=9) and anxiety (n=10), whereas none of the HC participants had depression and only 2 reported anxiety. All LC patients, but not HC, reported pain on the day of sampling, with an average intensity of 3.3 ± 2.84 at the Numeric Pain Rating Scale. Additionally, 8 LC patients were identified as suffering from neuropathic pain according to the DN4 questionnaire criteria. No significant differences in the serum biomarkers of neuronal or glial damages, respectively Nf-L, UCH-L1 and GFAP, were observed between the HC and LC groups (Table 2).

### IgG concentration in sera and purified fractions

IgG were purified from the serum of LC patients and HC subjects. In HC serum samples, the mean IgG concentration was 7.27 ± 1.2 mg/mL, while the IgG fraction following column G purification contained 6.74 ± 0.5 mg/mL. In LC serum samples, the mean IgG concentration was 7.30 ± 1.3 mg/mL, and 7.27 ± 1.2 mg/mL in the purified IgG fractions. These values indicate that only a minimal loss of IgG occurred during the purification process (Suppl.Fig.1). Other immunoglobulin isotypes (IgE, IgA, and IgM) were also quantified and showed a significant decrease following purification (Suppl.Fig.1). This confirms that the purified fractions were specifically enriched in IgG, with minimal contamination from other isotypes.

### Passive transfer of IgG from LC patients induces a transient mechanical allodynia

To investigate the potential pathogenicity of IgG from LC patients, purified IgG from LC patients or HC subjects were administered daily to C57Bl/6J mice by intraperitoneal injection (8mg/day) for four consecutive days. During the two weeks post-injections, pain-related behaviors were evaluated in mice using a battery of behavioral tests (Fig.1A).

First, thermal sensitivity was assessed by measuring paw withdrawal latency using the hot plate and Hargreaves tests. No significant change in thermal hypersensitivity was observed between HC and LC groups in both tests (Fig.1B-C). This effect remained unchanged even when the analysis was restricted to the subset of five LC patients who experienced heat- and cold-induced pain, according to the DN4 questionnaire and the Numeric Pain Rating Scale (NPRS) (Suppl. Fig.2A-B).

Second, mechanical allodynia was evaluated using the Von Frey filaments test. Passive transfer of IgG from LC patients induced a transient and significant decrease in the paw withdrawal threshold compared to HC (Fig.1D). This effect was still observed when narrowing the analysis to eight LC patients who specifically experienced hypersensitivity to mechanical pressure, as reported in their DN4 questionnaire and NPRS (Suppl. Fig.2C).

Third, we assessed general well-being of mice using the Nest-building score and Facial Grimace Scale (FGS). No significant differences were observed in nest building behavior between groups (Fig.1E). However, FGS score was significantly higher (FGS ≥ 1) in mice receiving IgG from LC patients on day 1, with no further differences detected at later time points (Fig.1F). When we restricted the analysis to the subset of LC patients who reported mechanical hypersensitivity, similar effects were observed (Suppl.Fig.2D-E). Abnormal FGS score was even extending until day 2 of the injection protocol (Suppl.Fig.2E).

### Pain behaviors are alleviated upon LC IgG enzymatic digestion or IgG depletion

To confirm that pain-related behaviors were mediated by IgG from LC patients, the transfer protocol was slightly adapted. In brief, IgG-depleted serum, papain-digested IgG or purified IgG fractions from the same batches of LC patients (n=3) were transferred to C57Bl/6J mice for two consecutive days. Pain-related behaviors were performed along this period (Fig.2A), as we previously demonstrated that mechanical allodynia was triggered from the first day of transfer.

Papain is a cysteine-protease cleaving human IgG at the hinge region, leading to a fragment crystallizable (Fc) and two fragment antigen-binding (Fab) domains. A western blot validated the effective enzymatic cleavage of the papain-digested fraction, compared to the total serum, native purified IgG, and IgG-depleted serum (Fig.2B). A band near 50 kDa, corresponding to the human IgG heavy chain, and a 25 kDa band for the light chain were detected in total and purified IgG samples. These bands were absent in IgG-depleted serum, confirming successful depletion. In the papain-digested samples, a new band at ±30 kDa (Fc fragment) and another at ±25 kDa (Fab fragment) were observed. However, the persistence of the 50 kDa heavy chain band indicated only partial digestion (Fig. 2B).

Thermal and mechanical hypersensitivities were abolished when mice were injected with IgG-depleted serum or papain-digested IgG, compared to those receiving native IgG from LC patients (Fig.2C-F). The thermal hypersensitivity was found to be more persistent over time in response to the cold plate than to the hot plate (Fig.2C-D).

### Passive transfer of human IgG from LC patients does not trigger cognitive deficits, anxiety, or depression

We next assessed spatial working memory using the Barnes maze (long-term memory) and Y-maze (short-term memory), anxiety-related behavior using the Elevated-plus maze and Light/Dark box tests, and depressive-like behavior using the Tail Suspension Test. In the Barnes maze, no significant difference was observed between groups in either the latency to locate the target hole (primary latency) and to enter into the escape box (total latency) (Fig.3A). Spontaneous alternation as well as total distance covered in the Y-maze did not differ between mice receiving IgG from LC patients and those receiving IgG from HC subjects (Fig.3B-C). Similar effects were observed when the analysis was restricted to mice receiving IgG from the four LC patients with cognitive impairments (Table 2), as defined by a MoCA score below 26 (Suppl.Fig.3A, B). In the same vein, the time spent in the closed arms of the Elevated Plus Maze, in the dark compartment of the Light/Dark box, as well as the immobility time at the Tail Suspension Test were all not different in mice that received IgG from this subset of LC patients (Fig.3D-E-F). Non-significant results were also obtained when the analysis was restricted to mice injected with IgG from the subset of patients with anxiety-only or depression-only outcomes (HADS score above 8) (Suppl.Fig.3C-E).

### IgG injections from LC patients do not induce microgliosis or astrogliosis

Pain-related behaviors induced in mice that received IgG from LC patients could be a consequence of neuroinflammation. Hence, we performed immunohistochemistry targeting Iba-1 to identify microglial cells and GFAP for astrocytes. We did not observe a significant difference in the density of Iba1-positive cells in different regions-of-interest of the brain, or in the lumbar spinal cord (Fig.4A-B). Similarly, there were no differences in GFAP-positive cells in the corpus callosum, hippocampus or brainstem and in the lumbar spinal cord (Fig.4D-E). These results were further supported by RT-qPCR analyses, which showed no significant changes of *Iba1* and *Gfap* mRNA expression in either whole-brain or DRG homogenates (Fig.4C,F).

### IgG from LC patients accumulates into the peripheral nervous system but not the central nervous system

Since mice injected with IgG from LC patients exhibited increased hypersensitivity, we investigated the distribution of human IgG within the nervous system to identify potential sites of binding or action. Immunohistochemical analyses were performed on tissues collected at the 4^th^ day of IgG injection protocol. No immunoreactivity for human IgG was found in the brain parenchyma and spinal cord (Fig.5A-B), except in the connective tissue and blood vessels located in the choroid plexuses and meninges of injected mice, in both transferred groups (Fig.5A). Respective tissues from naive mice, which did not undergo any transfer of human IgG, were devoid of immunoreactivity, and served as control for antibody species specificity. In contrast, human IgG were detected in the ganglionic cell body-rich area, mainly in the peri-neuronal spaces of dorsal root ganglia (DRG) of the HC group, while in the LC group, immunolabeling of neuron cell bodies was observed (arrows, Fig.5C), indicating a differential accumulation of IgG in peripheral sensory structures.

### IgG from LC patients binds to sensory neurons in the lumbar dorsal root ganglia

To identify the cell types targeted by human IgG in the DRG, we performed double immunofluorescence against neuron cell bodies (NeuN) and satellite glial cells (Glutamine Synthetase). As for immunohistochemistry, human IgG from HC subjects were detected in the peri-neuronal network of DRG. Human IgG from LC patients colocalized with NeuN, but not with glutamine synthetase (Fig.6A-C). The percentage of NeuN+/IgG+ cells was significantly higher in DRG from mice injected with LC patient IgG than in DRG transferred with HC IgG (Fig.6B). When characterizing DRG neuron subtypes using fiber-specific markers, we observed that approximately 40% of NF200^+^ neurons (Aβ and Aδ fibers) colocalized with human IgG, whereas about 18% of peripherin^+^ neurons (C-fibers) and 12% of cGRP^+^ neurons (C-fibers and Aδ fibers) showed IgG colocalization (Suppl.Fig.4). The colocalization between human IgG and sensory neurons was no longer observed when mice received IgG-depleted serum or papain-digested IgG compared to mice receiving purified LC IgG (Fig.7).

### Targeted antigen screening failed to identify a common autoreactivity profile across LC sera

We next sought to identify the putative antigens differentially recognized by IgG (auto)antibodies in LC patients. Serum samples were screened against a panel of 120 known neurological disease–associated autoantigens (Fig.8A). No striking difference was observed between the autoreactivity profiles of LC and HC groups, due to a high degree of inter-individual heterogeneity and the limited number of patient samples. Among the candidate autoantibodies close to the significance threshold, anti-HSPA5/GRP78 levels (*p*= 0.064) were found nearly doubled in the serum of LC patients, while NOVA1 and RBPJ antibodies (*p*= 0.057 and *p*=0.049, respectively) tend to be downregulated (Fig. 8B).

## Discussion

Although the underlying mechanisms behind long COVID remain unclear, growing evidence points towards a post-viral autoimmune syndrome, besides other concomitant drivers such as viral persistence. In this study, we specifically investigated the pathogenic effect of post-COVID circulating IgG and sought to identify their cellular targets within the nervous system. One approach to directly assess the causality is to perform passive transfer of purified human IgG to mice. When mice received IgG from LC patients, they exhibited increased mechanical sensitivity which resolved after cessation of IgG injections. This study strengthens (non-peer reviewed yet) data obtained on independent long COVID cohorts (USA and NL) [34, 35]. Several studies have shown that passive transfer of IgG from patients (fibromyalgia, complex regional pain syndrome) can recapitulate the pain symptomatology in mice [36–38]. Interaction between the nervous and immune system in the context of autoimmune diseases has been increasingly investigated in the pathophysiology of chronic pain conditions and nowadays referred as ‘autoimmune pain’. For instance, passive transfer of contactin-associated protein-like 2 (CASPR2) antibodies from neuropathic patients has been shown to trigger mechanical allodynia in mice [39]. Growing attention is also being paid to autoimmune diseases following viral infection. For instance, cases of rheumatoid arthritis are described following Epstein-Barr virus (EBV) infection [40]. A breakdown of self-tolerance and molecular mimicry are thought to play a central role, as anti-citrullinated protein antibodies (ACPA) directed against the EBV nuclear antigen-1 (EBNA-1) can cross-react with citrullinated fibrinogen in inflamed synovial tissue [41, 42].

In the context of long COVID pain-related symptoms, the exact antigenic target(s) still need(s) to be identified, knowing that a plethora of autoantibodies targeting nervous system epitopes have already been detected in the serum of patients [26–31]. Functional investigations of these autoantibodies still need to determine whether they are pathogenic, or accessory antibodies devoid of clinical significance. Some research outcomes demonstrated that LC patients have functional anti-GPCR antibodies (anti-β_2_ adrenoceptor, muscarinic M2 receptor, α_1_ -adrenoceptor, endothelin receptor, nociceptin receptor) [27, 43]. Moreover, the serum concentration of these autoantibodies is correlated to disease severity [26, 28, 44]. Interestingly, the combination of β_2_ adrenoceptor and muscarinic M2 receptor antibodies was previously identified in patients with complex regional pain syndrome [45].

Our findings further support the involvement of IgG in the symptomatology of LC patients, as well as their structural integrity as demonstrated by the alleviation of pain phenotype when mice were transferred with papain-digested IgG. The same relief effect was observed when mice received IgG-depleted serum, which mimics, from a translational point-of-view, immunoglobulin apheresis. Clinical improvement has been reported following therapeutic apheresis [32] or treatment with intravenous immunoglobulin (IVIG) [46, 47]. As such, IVIG can neutralize autoantibodies, saturate neonatal fragment crystallizable receptors (FcRNs) and inhibit subsequent complement activation [46]. Ongoing clinical trials (NCT05350774; NCT06305793; NCT06524739) are evaluating IVIG efficacy on LC patients. Thompson and others reported that on nine LC patients included with immune dysregulation, six were treated with IVIG for three months and they experienced both symptomatic and laboratory improvements along with the ability to return to work [47]. Similarly, sixteen LC patients with small fiber neuropathy symptoms, of whom nine were treated with IVIG 17 months post-acute infection. At six months post-treatment, all patients showed symptoms improvement, with six presenting complete resolution and three with substantial improvement of neuropathic symptoms [48]. Altogether, these observations strongly suggest that treatments targeting autoantibodies can alleviate pain-related symptoms.

Brain fog, attention, and concentration deficits accounting among the most prevalent symptoms in LC patients, we investigated whether passive transfer of IgG mimicked cognitive impairments in mice. Our results showed that mice did not experience memory impairment, anxiety-, depressive-like behaviors or fatigue. Immunohistochemistry against human IgG failed to detect IgG in memory- or cognition-associated brain regions (hippocampus and cortex). Finally, no signs of brain inflammation were observed in mice that received an injection of IgG from patients with long COVID. Thus, either IgGs do not play a role in the cognitive impairment of long COVID patients or our mouse model cannot fully recapitulate the human pathology. For example, human IgG might not recognize murine epitopes or might not cross an intact blood-brain barrier.

Unlike brain or spinal cord parenchyma, transferred human IgG were found in murine lumbar dorsal root ganglia (DRG) with distinct patterns: in the peri-neuronal spaces in the HC group and on neuronal cell bodies in the LC group. The peri-neuronal distribution observed in controls is not unexpected, as DRG are highly vascularized structures endowed with fenestrated capillaries that allow permeability to both low and high molecular weight molecules [49]. IgG from LC patients colocalized with neuron cell bodies but not with satellite glial cells (SGC). These findings differ from other preprints reporting IgG binding to murine DRG with no group differences, [34, 35]. In contrast, IgG from fibromyalgia patients primarily target SGC [36, 38] while in small fiber neuropathy, anti-plexin D1 antibodies specifically bind to nociceptive neurons [50]. Passive transfer of these antibodies to mice induced mechanical and thermal sensitivity [36, 38, 50]. More specifically in our study, IgG from LC patients preferentially targeted subsets of sensory neurons, particularly A-fibers and, to a lesser extent, C-fibers. A-fibers can be divided into two main categories: Aβ-fibers, which are involved in proprioception and Aδ-fibers which mediate mechanoreception and nociception. C-fibers participate in nociception and thermoception. This dual involvement in both A- and C-fiber populations has already been reported in the context of small fiber neuropathy [51]. The pronociceptive effects of IgG from LC patients may involve immune complexes interacting with Fc gamma receptors (FcγR) expressed by sensory neurons. Such interaction can modulate nociceptive signaling by modulating neuronal excitability [52]. Consistently, mechanical allodynia and DRG hyperexcitability were no longer present in nerve-injured FcγR-deficient mice [53]. Although binding of IgG-immune complexes to FcγR may have promoted inflammation [52], neither astrocytic nor microglial activation were evidenced in DRG tissues, brain and spinal cord in our model of passive transfer.

A major challenge in LC research is differentiating newly developed post-infectious symptoms from pre-existing, subclinical, or previously undiagnosed conditions that only became apparent after infection. The present study bears limitations, the first one being the limited number of LC patients and the overrepresentation of female individuals in the cohort compared to overall LC prevalence. However, the patients were carefully selected to ensure a high degree of homogeneity, thus strengthening the observed results. Second, although passive transfer models are valuable tools to demonstrate the pathogenic potential of antibodies, they typically rely on short-term, repeated high-dose administrations of IgG. As such, they may not fully reproduce the prolonged and complex immune interactions occurring in patients with a chronic condition. In this paradigm, it should be kept in mind that human IgG may not – fully or at all - recognize murine epitopes or efficiently bind to the murine Fc receptor that could mediate downstream pathogenic events. In addition, heterogeneity of clinical manifestations suggests that only certain autoantibodies are present in subsets of patients and capable of inducing symptoms in mice, while others have no effect. Also, our study only used female mice as LC patients are mainly women, and it is well-known that pain and autoimmune mechanisms may involve sex-specific pathways that may not be fully recapitulated in male mice [54]. Finally, our antigen microarray failed to identify significantly upregulated levels of autoantibodies; because it used a biased approach toward 120 known nervous system-derived autoantigens and thus may have missed relevant genuine targets. Interestingly, data from a preprint, based on the screening of more than 21,000 human antigens, suggest unusually high levels of autoantibodies targeting the MED20 and USP5 proteins in the serum of LC patients; however, no specific causal relationship with the disease phenotype has yet been demonstrated [35]. Further studies are needed to identify the specific autoantibody(ies) using unbiased autoantibodyome screening, as well as their mechanisms-of-action towards DRG structures or related nerve endings. In that vein, complementary experiments using *in vitro* approaches, based i.e. on human iPS-derived sensory neurons, might focus on a potential antibody-dependent cellular cytotoxicity or a modulation of their intrinsic excitability.

## Conclusion

In conclusion, our data comfort the hypothesis of autoimmunity underlying pain-related symptoms in LC patients while providing new insights on the target of these IgG which are sensory neurons of lumbar dorsal root ganglia (DRG). Future research should focus on investigating the antigenic target and mechanisms behind DRG sensory neurons dysfunction.

## Acknowledgements

Margaux Mignolet is supported by a FNRS/FRIA doctoral fellowship. This research program benefited from fundings granted to Charles Nicaise (FNRS CDR J.0147.22). We thank Dr. Dominique Butenda Babapu (CHU-UCL Namur) for referring LC patients and Dr. Caroline Meyer (CHU-UCL Namur) for providing the pain-related questionnaires. We acknowledge Dr. Benoît Vokaer who instilled the seminal idea of IgG-mediated pain symptoms. This research was made possible thanks to access to the microscope facility of the “Plateforme Technologique Morphologie – Imagerie” and the “Plateforme Technologique Sciences de la Vie” (Université de Namur). We would like to thank all the participants to the study for their contribution.

## List of abbreviations

CASPR2: contactin-associated protein-like 2
COVID-19: coronavirus-associated disease 2019
DRG: Dorsal Root Ganglia
EBV: Epstein-Barr virus
FGS: Facial Grimace Scale
GFAP: Glial Fibrillary acidic protein
HADS: Hospital Anxiety and Depression Scale
Iba1: Ionized calcium-binding adapter molecule 1
IVIG: Intravenous Immunoglobulin
HC: Healthy Control
LC: Long COVID
MoCA: Montreal Cognitive Assessment
Nf-l: Neurofilament-light
NPRS: Numeric Pain Rating Scale
PASC: post-acute sequelae of COVID
SARS-CoV-2: severe acute respiratory syndrome coronavirus 2
SDMT: Symbol Digit Modalities Test
SGC: Satellite Glial Cells
UCH-L1: Ubiquitin carboxy-terminal hydrolase L1
WHO: World Health Organization

## Declarations

### Ethics declarations

#### Ethics approval and consent to participate

Human ethic project was approved by the CHU-UCL Namur ethics Committee (157.2022). Written and informed consent were obtained for all study participants. The experimental procedure was approved by the Animal Ethics Committee of the University of Namur (ethic project UN 23-392 and UN 24-438).

#### Consent for publication

All co-authors approved the latest version of this manuscript and provided their consent for submission.

#### Authors’ contributions

Conceptualization: CN, NG, PB, MM ; Methology M.M., V.B., K.D.S, T.F.; software, M.M., T.F.; validation, M.M., C.N.; formal analysis M.M., T.F.; investigation M.M., T.F.; resources V.B., K.D.S., C.D., F.G., M.J., P.B.; data curation M.M., T.F.; writing – original draft preparation M.M., C.N.; writing-review and editing M.M., N.G., J.G., P.B., C.N.; visualization M.M., C.N.; supervision C.N., N.G., P.B.; projection administration C.N.; funding acquisition C.N.

