## Supplemental figures for "Pathogenic IgG from long COVID patients with neurological sequelae triggers sensitive but not cognitive impairments upon transfer into mice"

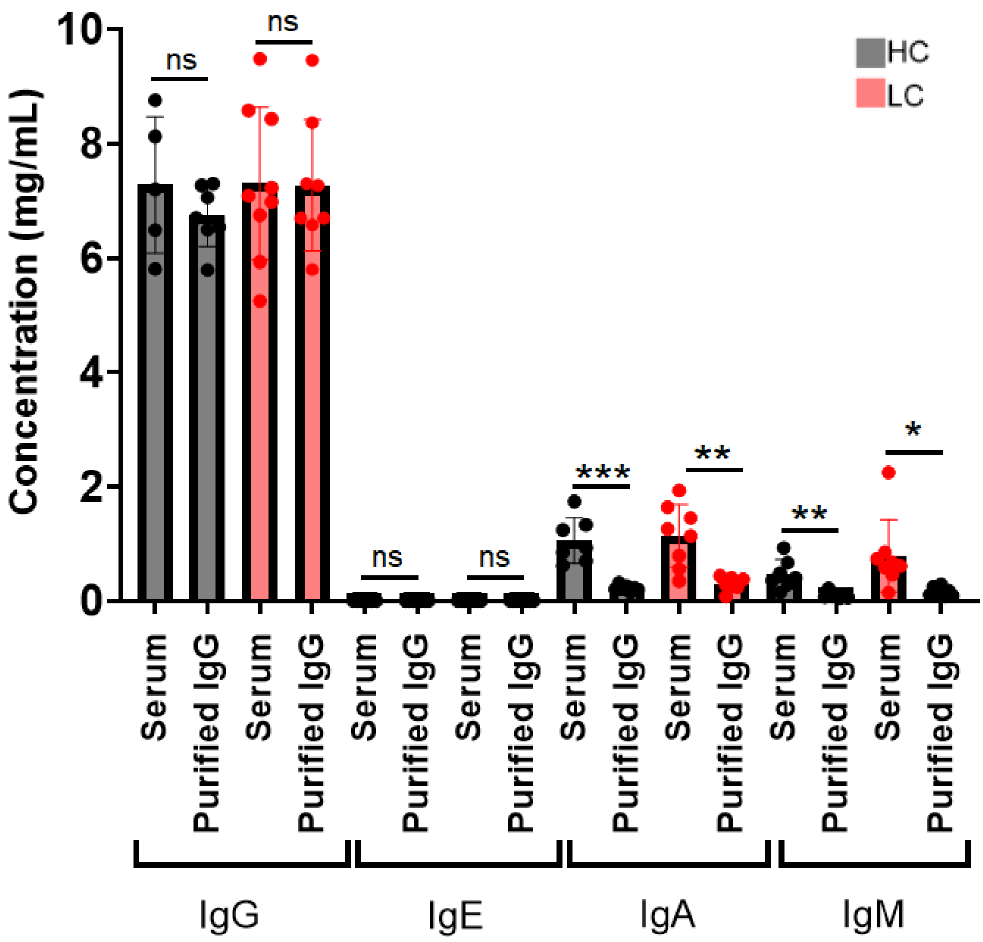


**Supp.Fig.1. Characterization of immunoglobulin isotypes in the serum and purified IgG fractions.** n=8 healthy controls (black box) and n=7 long COVID patients (red box). IgG, IgE, IgA, IgM isotypes were all quantified in the serum before and after column G purification and showed a specific enrichment of IgG in purified fractions. Each data point is the measured immunoglobulins for one HC or LC patient. Mean ± SD. Statistical analyses were computed using a paired t test *(*p<0.05, **p<0.01, ***p<0.001).*

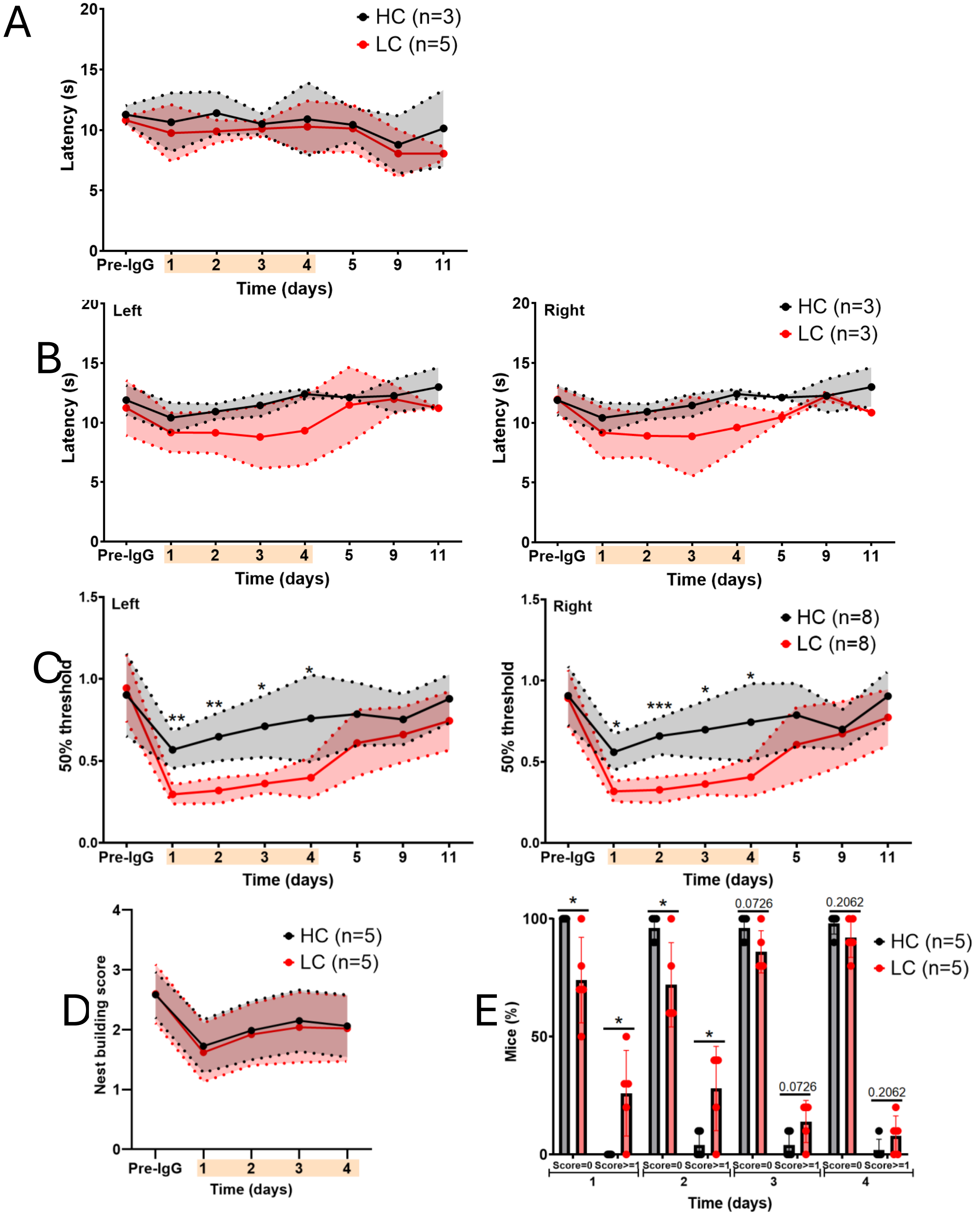


**Suppl.Fig.2. Pain-related behavioral tests for mice injected with IgG from long COVID patients with a DN4 score ≥ 4.** The same analysis as in Figure 1 was carried out considering only the subset of patients with a score of 4 or more at the DN4 questionnaire assessing patient-reported criteria of neuropathic pain. n=3-5 HC and n=3-8 LC. 10 mice were used per patient IgG batch. The orange overlay corresponds to the injection period. (A) Paw withdrawal latency at the hot plate test was unchanged between HC and LC groups. (B) Hind paw (left and right) withdrawal latency at the Hargreaves test did not differ significantly between HC and LC groups. (C) Hind paw (left and right) withdrawal threshold at the Von Frey filaments was significantly decreased during the first four days post-injection in LC condition. (E) The general well-being and motivation behavior were similar between groups of mice at the nest building score. (F) The percentage of mice with an abnormal Facial Grimace Scale (score ≥1) was significantly increased along the two first days of injections in LC group. Mean ± SD. Mixed-effects model followed by a Holm-Sidak multiple comparison test between HC and LC *(*p<0.05, **p<0.01, ***p<0.001).*

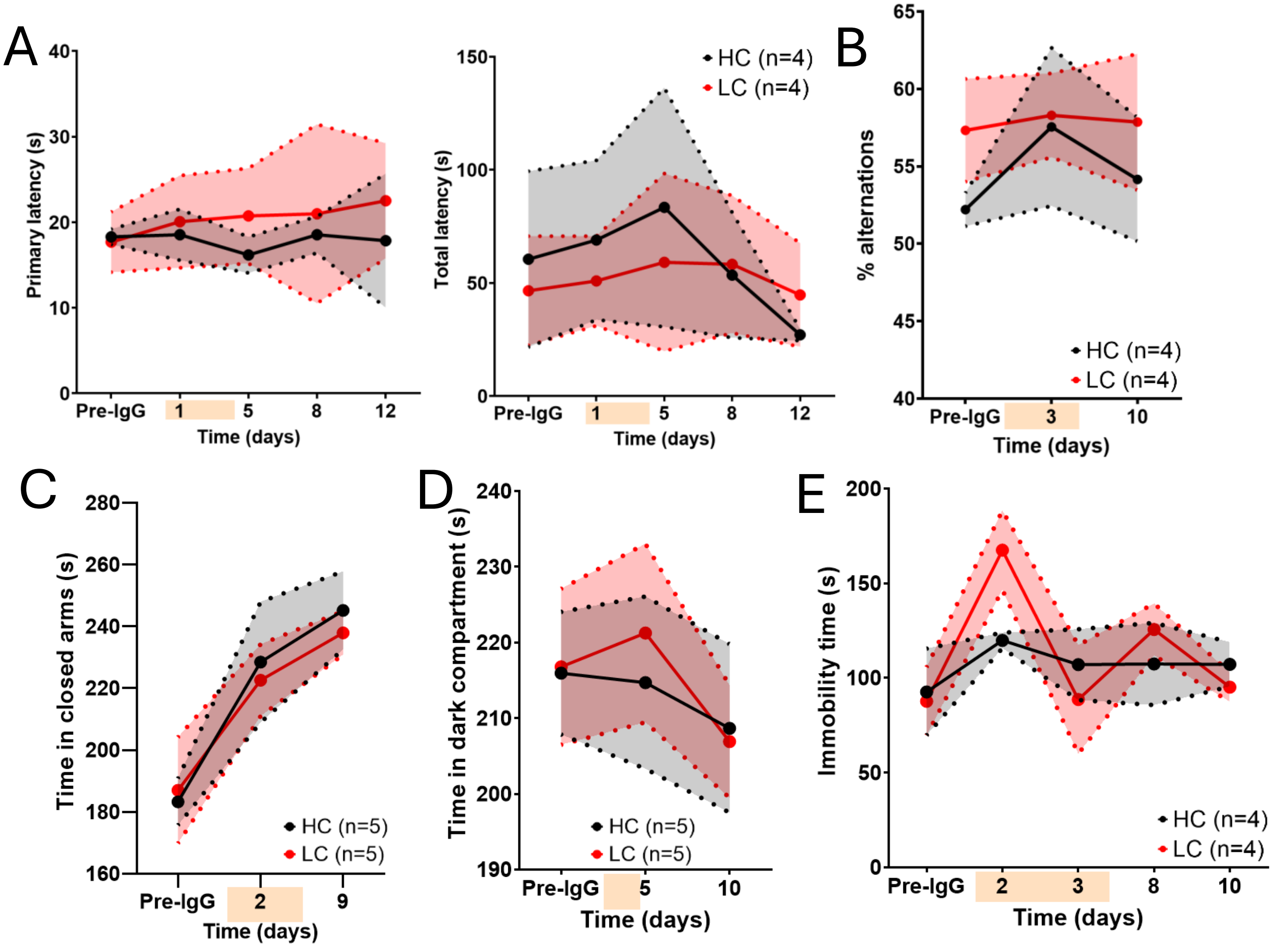


**Suppl.Fig.3. Spatial memory-, anxiety- and depression-related behavioral tests in mice transferred with IgG from subsets of patients with a Moca score ≤ 26 or HAD score ≥ 8.** The same analysis as in Figure 3 was conducted considering only the subset of patients with a Moca score below or equal 26 or anxiety/depression score above or equal 8. n=4-5 HC and n=4-5 LC patients. 10 mice were used per patient IgG batch. The orange overlay corresponds to the injection period. (A) Primary and total latencies at the Barnes maze did not differ between HC and LC groups. (B) Alternation between the arms of the Y-maze did not differ between HC and LC groups. (C) Time spent in the closed arms of the elevated-plus maze, as a measure of anxiety, was similar between groups. (D) Time spent in the dark compartment of the light and dark box, as a measure of anxiety, was similar between groups. (E) Immobility time at the tail suspension test, as a proxy of depressive-like behavior in mice, did not differ between HC and LC groups. Mean ± SD. Mixed effects model followed by a Holm-Sidak multiple comparison test between HC and LC.

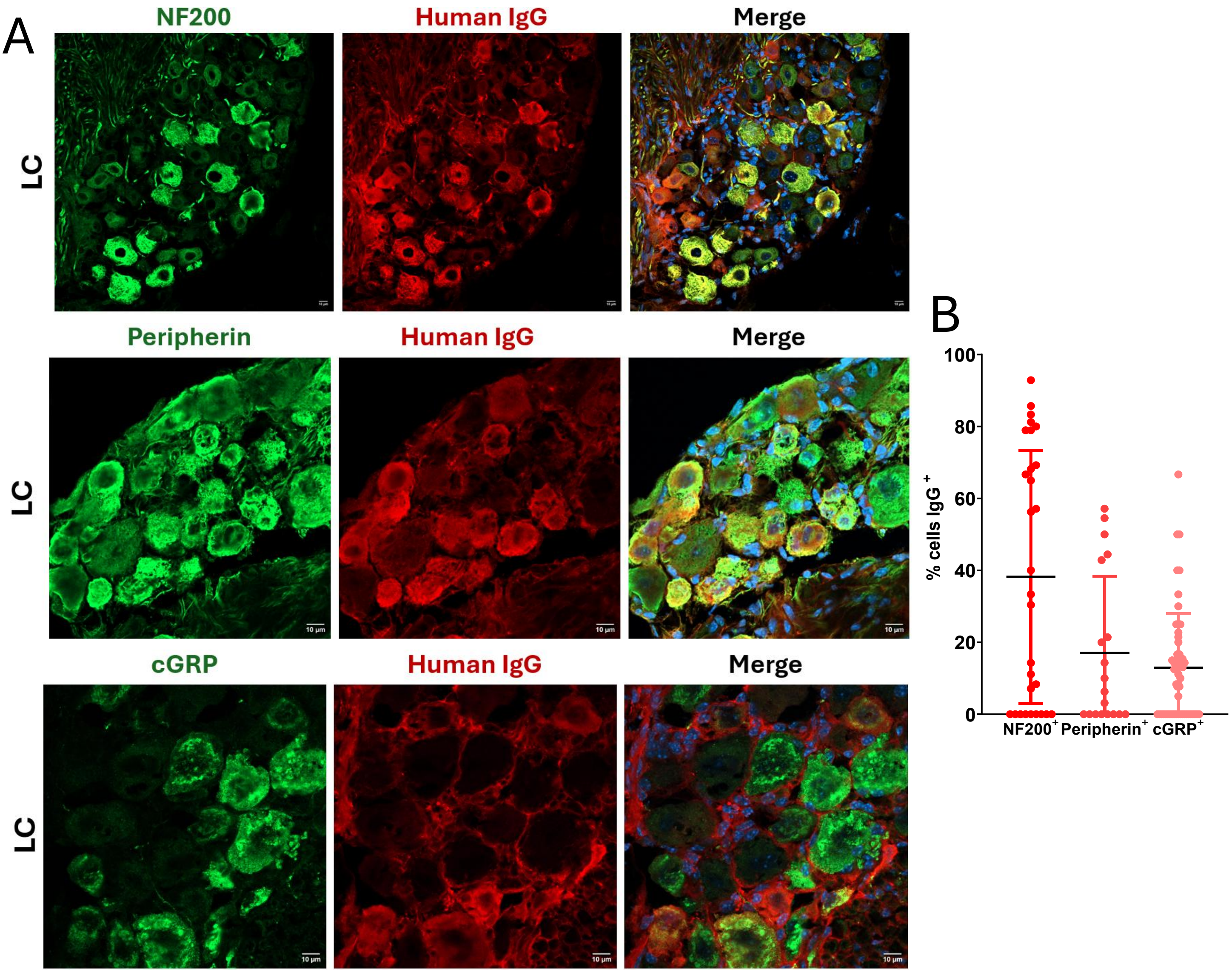


**Suppl.Fig.4. Colocalization of human IgG with sensory neurons subtypes in the mouse DRG after transfer of IgG from long COVID patients (LC).** Human IgG were immunolabeled using fluorescent anti-human IgG antibody (red). Nuclei were counterstained with Hoechst (blue). (A) Neuronal cell bodies in the murine DRG were immunostained (green) with anti-NF200 (A-fibers) or anti-peripherin (C-fibers) or anti-cGRP (peptidergic or Aδ-fiber neurons) antibodies. (B) Quantification of $NF200^+/IgG^+$ or $peripherin^+/IgG^+$ or $cGRP^+/IgG^+$ cells was performed using FIJI ImageJ. Each dot represents a colocalization. DRG included in the quantitative analysis were from 3 LC (n=6 mice). Mean ± SD. Scale bar represents 10 µm.
